# Structural elucidation and antiviral activity of cathepsin L inhibitors with carbonyl and epoxide warheads

**DOI:** 10.1101/2023.08.11.552671

**Authors:** Sven Falke, Julia Lieske, Alexander Herrmann, Jure Loboda, Sebastian Günther, Patrick YA Reinke, Wiebke Ewert, Katarina Karničar, Aleksandra Usenik, Nataša Lindič, Andreja Sekirnik, Hideaki Tsuge, Vito Turk, Henry N Chapman, Winfried Hinrichs, Gregor Ebert, Dušan Turk, Alke Meents

**Affiliations:** Center for Free-Electron Laser Science CFEL, Deutsches Elektronen-Synchrotron DESY, Notkestraße 85, 22607 Hamburg, Germany; Institute of Virology, Helmholtz Munich, Trogerstrasse 30, 81675 Munich, Germany; Department of Biochemistry and Molecular and Structural Biology, Jozef Stefan Institute, Jamova 39, 1000 Ljubljana, Slovenia; Centre of Excellence for Integrated Approaches in Chemistry and Biology of Proteins, Jamova 39, 1000 Ljubljana, Slovenia; Faculty of Life Sciences, Kyoto Sangyo University, Kyoto 603-8555, Japan; Hamburg Centre for Ultrafast Imaging, Universität Hamburg, Luruper Chaussee 149, 22761 Hamburg, Germany; Department of Physics, Universität Hamburg, Luruper Chaussee 149, 22761 Hamburg, Germany; Institute of Biochemistry, Universität Greifswald, Felix-Hausdorff-Str. 4, 17489 Greifswald, Germany

**Keywords:** Cathepsin L, SARS-CoV-2, crystallography, active site cysteine modification, drug development, dual targeting, virus entry, anti-viral compound

## Abstract

Emerging RNA viruses including SARS-CoV-2 continue to be a major threat around the globe. The cell entry of SARS-CoV-2 particles via the endosomal pathway involves the cysteine protease cathepsin L (CatL) among other proteases. CatL is rendered as a promising drug target in the context of different viral and lysosome-related diseases. Hence, drug discovery and structure-based optimization of inhibitors is of high pharmaceutical interest. We herein verified and compared the anti-SARS-CoV-2 activity of a set of carbonyl and succinyl-epoxide-based inhibitors, which have previously been identified as cathepsin inhibitors. Calpain inhibitor XII (CI-XII), MG-101 and CatL inhibitor IV (CLI-IV) possess antiviral activity in the very low nanomolar IC_50_ range in Vero E6 cells. Experimental structural data on how these and related compounds bind to CatL are however notably lacking, despite their therapeutic potential. Consequently, we present and compare crystal structures of CatL in complex with 14 compounds, namely BOCA (*N*-BOC-2-aminoacetaldehyde), CLI-IV, CI-III, CI-VI, CI-XII, the main protease α-ketoamide inhibitor 13b, MG-101, MG-132 as well as E-64d (aloxistatin), E-64, CLIK148, CAA0225, TC-I (CID 16725315) and TPCK at resolutions better than 2 Å. Overall, the presented data comprise a broad and solid basis for structure-guided understanding and optimization of CatL inhibitors towards protease drug development.

## 1. Introduction

In addition to other emerging RNA viruses, Betacoronaviruses remain to be a major global health concern. More than six million cumulated deaths following a *severe acute respiratory syndrome coronavirus 2* (SARS-CoV-2) infection were reported for the time between March 2020 and the end of March 2022 (*WHO COVID-19 dashboard;* https://covid19.who.int/*; last access: 07.06.2023*). It is well established that human cathepsins and in particular the lysosomal cysteine protease cathepsin L (CatL) are involved in the cell entry of SARS-CoV and SARS-CoV-2 via endosomes^1^ – a path alternative to the viral cell entry utilizing the serine protease TMPRSS2.^2^ CatL can proteolytically process the surface exposed trimeric spike protein of SARS-CoV-2 particles, which then enter the cell via clathrin-coated vesicles.^3,4^ The Omicron variant of SARS-CoV-2 appears to utilize this endosomal entry pathway even more efficiently compared to previously originating variants of the virus.^5^ Hence, CatL was specifically identified as an attractive host-cell drug target to interfere with COVID-19.^6^ Beside coronaviruses, CatL has additional relevance as a drug target as it is involved in activation of Hendra virus and Nipah virus fusion protein and thereby required for replication.^7,8^ CatL has further been reported as an important drug target for the treatment of nuclear lamina damage in Alzheimer’s disease^9^, cancer^10^ and other diseases.^11–13^

CatL is nearly ubiquitously expressed.^13^ *In vivo*, CatL has multiple functions and is active on a variety of substrates at slightly alkaline and over a broad range of acidic pH values with an optimum of 5.5 for elastin.^14^ It prefers combinations of hydrophobic residues at P3 and P2 positions. Several amino acid combinations favor positively charged residues at P1 and P1’ positions.^15,16^ The common Schechter and Berger nomenclature of S and P subsites of protease active sites^17^ will be used consistently herein.

Proteases are generally considered as attractive drug targets due to their essential signaling roles in activation of other enzymes. Given the chemical diversity of protease inhibitors already available, they may be used as a starting point to adjust it to another target^18^. Basic covalent inhibition of cysteine proteases can be achieved – among other functional groups – via a vinylsulfone^19^, a halomethyl ketone^20^, an epoxide^21^ an aldehyde^22^, a ketoamide moiety^23^ or an alkyne^24^ reacting with the nucleophilic thiolate group of the active site cysteine. Inhibition is frequently supported by a peptidomimetic scaffold binding to the substrate recognition site. Peptides with halomethyl ketone warhead are additionally well known to inhibit serine proteases by specific covalent binding to the active site serine and also to the catalytic histidine.^25^

Interestingly, many cathepsin-targeting protease inhibitors were discovered and developed based on an activity screening. Although *in vitro* assays for a number of CatL inhibitors are available and provide IC_50_ or even more valuable K_i_ values due to the complex binding mechanism^6,26–29^, structural data on how they bind to CatL are scarce. Therefore, it was the goal of the present work to test different protease inhibitors for their activity against SARS-CoV-2 and to structurally elucidate and understand their mode of action on the atomic level.

For our work, different protease inhibitors with either reported SARS-CoV-2 anti-viral activity and/or known cathepsin inhibition were selected (Table 1). The calpain inhibitor calpeptin, which was initially identified as an anti-SARS-CoV-2 drug targeting M^pro^, and more recently identified as a highly potent cathepsin inhibitor suggesting a so-called dual targeting approach of both SARS-CoV-2 M^pro^ and CatL^30–32^, was included as a reference. Likewise, calpain inhibitor XII (CI-XII) has been reported as another antiviral dual-target inhibitor.^33^ MG-132, CAA0225, and TC-I (CID 16725315) have further been reported to have anti-coronaviral activity.^27,30,34,35^ Parameters of CatL inhibition by these compounds and further references are provided in table S1. The compounds contain different warheads expected to bind to the active site cysteine of CatL.

**Table 1.**
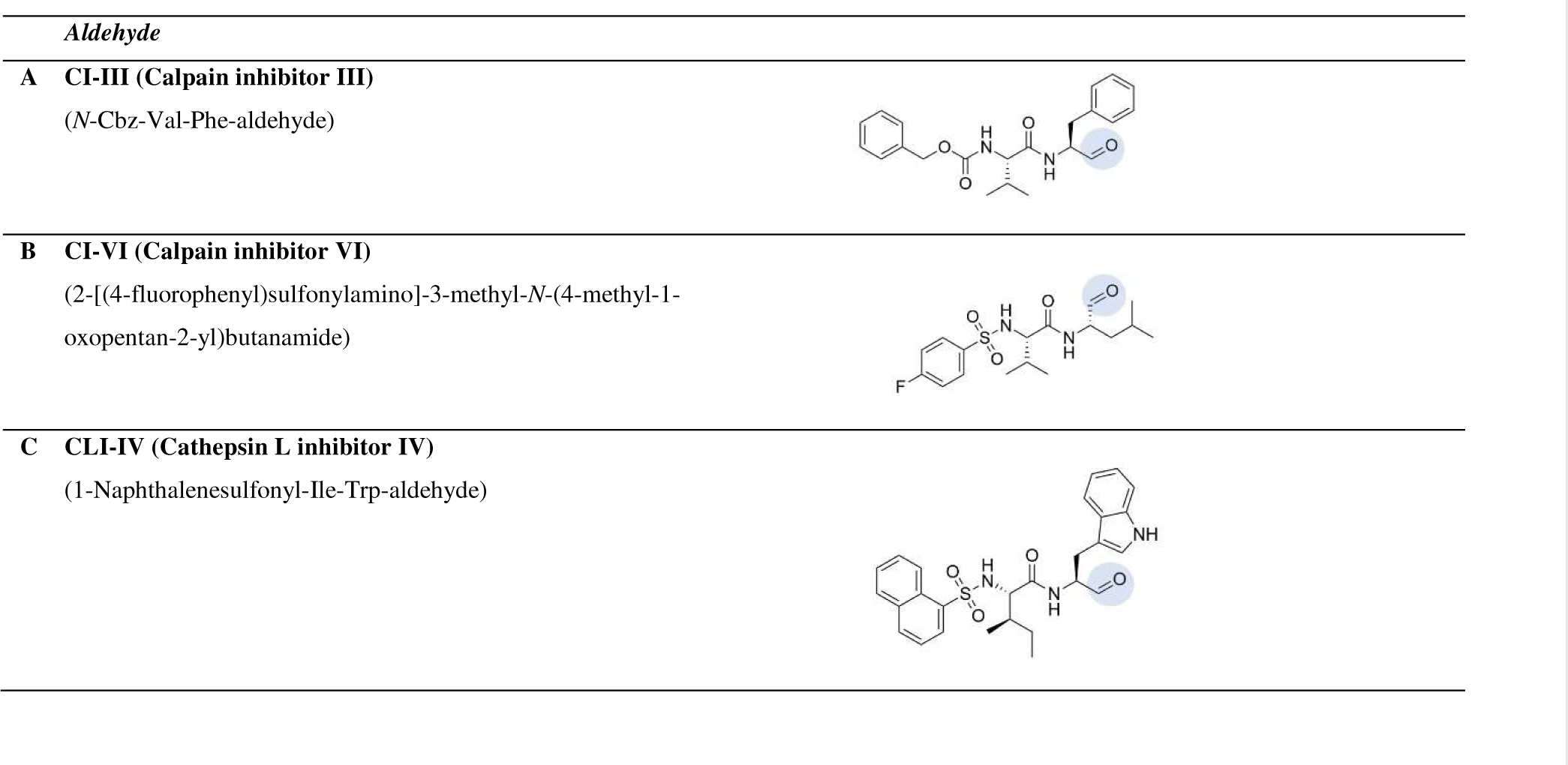
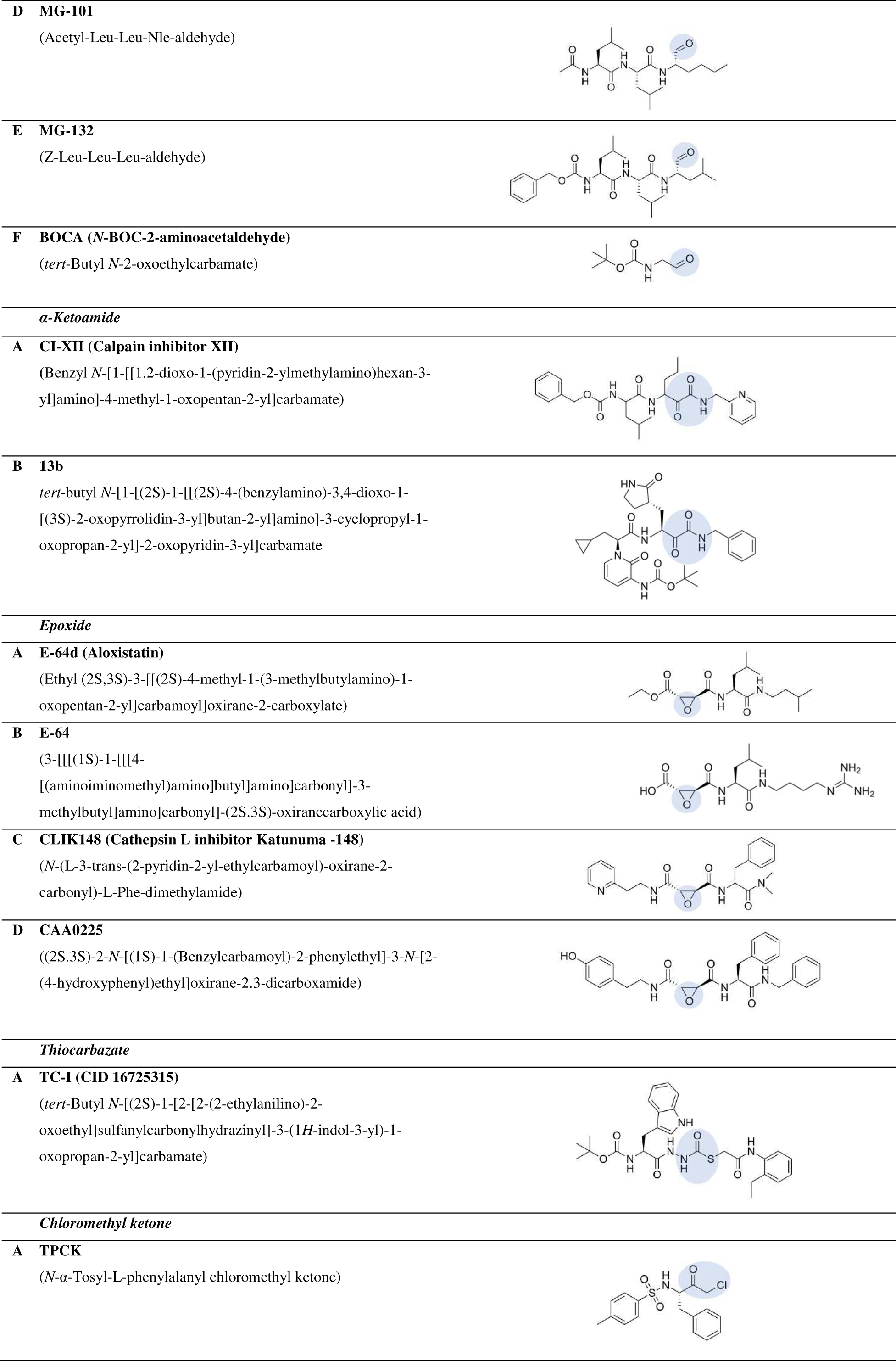
Cathepsin inhibitors under investigation grouped according to the respective warhead. The reactive site is highlighted in light blue. Covalent binding is schematically illustrated in figure S1.

Seven inhibitors showed distinct anti-viral activity against SARS-CoV-2 as determined in Vero-E6 cells using a fluorescence detection principle. A high anti-viral potency with IC_50_ values in the low nanomolar range was observed for CI-XII, MG-101 and cathepsin L inhibitor IV (CLI-IV). Most importantly, to complement the data and elucidate the inhibition, X-ray crystal structures of the 14 compounds listed in table 1 in complex with CatL were determined at resolutions better than 2 Å. This includes that we elucidated the interaction of CatL with 13b, a potent α-ketoamide drug, which has been reported to covalently inhibit the main protease of Alpha- and Betacoronaviruses – including SARS-CoV-2 M^pro^ – as well as the 3C protease of Enteroviruses.^23,36^

The presented high-resolution structural data, as well as activity assessment and nanoDSF data provide an experimental basis for detailed structure-based optimization of CatL drugs. The data particularly suggest to consider the scaffolds of CI-XII and MG-101 due to their dual-targeting and high anti-viral potency for further cathepsin drug development.

## 2. Materials and Methods

### 2.1. SARS-CoV-2 replication inhibition assay

Vero E6 cells have been used for viral growth and infection assays under culturing conditions that were previously described by Stukalov *et al.*^37^ The cell line was tested to be mycoplasma-free.

For virus production, Vero E6 cells (in DMEM, 5% FCS, 100 µg ml^-1^ streptomycin, 100 IU ml^-1^ penicillin) have been inoculated at a multiplicity of infection (MOI) of 0.05 with virus stock of SARS-CoV-2-GFP strain^38^ or SARS-CoV-2 Omicron strain BA.1. After 60 h of incubation at cell culture conditions (37 °C, 5% CO_2_), virus-containing supernatant was harvested, spun twice (1000 × g for 10 min) and stored at −80 °C. Viral titers were determined by performing a plaque assay. 20,000 Vero E6 cells per well were seeded 24 h before inoculation with five-fold serial dilutions of untitered virus stock and incubated for 1 h at 37 °C. After the designated incubation time, virus inoculum was exchanged with serum-free MEM (Gibco, Life Technologies) containing 0.75% carboxymethylcellulose (Sigma-Aldrich, high viscosity grade) and incubated for 48 h (37 °C, 5% CO_2_). Hereafter, cells were fixed with 4% PFA (20 min at RT), washed extensively with PBS before staining with 1% crystal violet and 10% ethanol in H_2_O for 20 min at RT, another washing step and finally calculating virus titers by counting of plaques.

After Vero E6 cells have been seeded and incubated overnight (10,000 cells per well in 96-well plates or 50,000 cells per well in 24-well plates), cells were treated with different concentrations of the inhibitors for 1 h before inoculation with SARS-CoV-2-GFP or SARS-CoV-2 (Omicron strain BA.1) at an MOI of 0.05.

As read-outs, either quantitative analysis of relative levels of SARS-CoV-2 qRT-PCR (non-GFP virus strain) or live-cell imaging (SARS-CoV-2 GFP) were performed. Live-cell imaging was conducted with an EssenBioscience IncuCyte with IncuCyte 2020C Rev1 software, taking pictures every 3 h (scan type: standard; image channels: Phase, Green to detect GFP; objective: 4×). Integrated intensity of detected signal in the Green channel was calculated by the IncuCyte 2020C Rev1 software.

For qRT-PCR, cells were harvested 24 h post inoculation. RNA was extracted using a NucleoSpin RNA kit (Macherey Nagel), according to the manufacturer’s protocol, eluting RNA in a volume of 50 µl. To transcribe 1 µl of yielded RNA into cDNA, PrimeScriptTM RT Master Mix (TaKaRa) was used according to manufacturer’s recommendations. Quantitative PCR was performed with the QuantStudio 3 system (ThermoFisher Scientific), using PowerUpTM SYBRTM Green Mastermix (AppliedBiosystems) to detect SARS-CoV-2 N transcripts (forward primer: TTA CAA ACA TTG GCC GCA AA; reverse primer: GCG CGA CAT TCC GAA GAA). Primers for RLPL0 transcripts were used as internal reference gene (forward, GGATCTGCTGCATCTGCTTG; reverse, GCGACCTGGAAGTCCAACTA). Data was analyzed with the second derivative maximum method. The relative amount of SARS-CoV-2 N transcripts in treated versus untreated cells was calculated by the 2(−ΔΔCt) method, using RLPL0 as a reference gene.

### 2.2. Production and purification of CatL

Gene overexpression of human procathepsin L was performed using *Pichia pastoris* strain GS115 (Invitrogen) based on a previously described procedure, along with the purification.^39^ The procathepsin L was purified via a pre-packed Ni-NTA affinity chromatography column and subsequent size-exclusion chromatography. Procathepsin L was auto-activated at 37 °C for approximately 3 hours. The sample was then applied to cation exchange chromatography resin SP Sepharose Fast Flow (Cytiva). The purified, activated CatL was reversibly blocked by a 10-fold molar amount of *S*-methyl-methanethiosulfonate and stored at −80 °C until further use. The dispersity of the protein solution was verified using dynamic light scattering. A Spectrolight 600 instrument (XtalConcepts) with a 660 nm red-light laser was used with 2 µl sample solution provided in a Terazaki plate covered by paraffin oil at room temperature. In preparation, the protein was dissolved in 50 mM sodium acetate, 100 mM NaCl, 1 mM TCEP, 500 µM EDTA, adjusted to pH 5.0, at a concentration of 40 µM.

### 2.3. NanoDSF

Nano Differential Scanning Fluorimetry (nanoDSF) measurements were performed with a Prometheus NT.48 fluorimeter (Nanotemper) using Prometheus Premium grade capillaries (Nanotemper). The excitation power was adjusted to obtain fluorescence counts above 2,000 RFU for 330 and 350 nm wavelength. The stability of CatL was investigated following the fluorescence ratio for the two wavelengths (F330/F350) depending on the solution temperature. For all compound measurements a final CatL concentration of 5 µM in 50 mM sodium acetate, 100 mM NaCl, 1 mM tris(2-carboxyethyl)phosphine (TCEP), 500 µM EDTA at pH 5.0 containing 2% (v/v) DMSO was used. For a melting temperature-based affinity screening a two-fold and 20-fold molar amount of the compound were used. Compound stock solutions were prepared in DMSO. SARS-CoV-2 M^pro^ was purified as described previously^30^ and a final protein concentration of 8 µM in 25 mM Tris, 100 mM NaCl, 1 mM TCEP adjusted to pH 7.5 and supplemented with 2% (v/v) DMSO was prepared. After incubation for 30 min at room temperature, the solutions, which were prepared in duplicates, were transferred to capillaries that were subsequently placed inside the fluorimeter. Data analysis was partly based on customized python scripts and the publicly available eSPC data analysis platform.^40^

### 2.4. Crystallization

Activated CatL (see section 2.2) concentrated to 7 mg ml^-1^ was equilibrated against 27% w/v PEG 8000, 1 mM TCEP and 0.1 M sodium acetate at pH 4.0 by sitting drop vapor diffusion in MRC maxi plates. Crystals, which grew to final size after approximately 3 days at 20 °C, were transferred to a soaking solution containing 22% w/v PEG 8000, 1 mM TCEP and 0.1 M sodium acetate at pH 4.0 as well as 5% v/v DMSO and 10% v/v PEG 400 for cryoprotection. In this solution crystals were soaked with selected compounds at approximately 1 mM final concentration for 24 h at 20 °C.

### 2.5. Diffraction data collection, processing and refinement

Crystals manually harvested in mother liquor soaking solution with PEG 400 were flash frozen in liquid nitrogen. Diffraction data were collected at 100 K at beamline P11 of the PETRA III storage ring (DESY, Germany) and subsequently processed with XDS.^41^ To reach optimal completeness, three data sets recorded at different positions of the same crystal were merged and scaled either with XSCALE^41^ or pointless/aimless.^42^ Initial atom coordinates were obtained by molecular replacement using Phaser^43^ and PDB 3OF9 as search model. Coordinates were iteratively refined using Phenix^44^ and via manual model building in Coot.^45^ Data collection and refinement statistics are provided in tables S2-4. Additional structure analysis and visualization was done using PyMOL (Schrödinger), Discovery Studio Visualizer (Biovia) and Poseview.^46^

### 2.6. Inactivation assay probing the inhibitors CAA0225, E-64 and CLIK148

CatL activity at a concentration of 5 nM in the presence of seven different concentrations of the inhibitors and in the absence of inhibitor (control sample) was monitored in real time for 30 min at 37 °C in 96-well black flat bottom microplates (Greiner) using an INFINITE M1000 pro plate reader (Tecan), the fluorescent peptide substrate Z-RR-AMC, and excitation and emission wavelengths of 370 and 460 nm, respectively. Inactivation rates were determined in two different buffers: 50 mM sodium phosphate buffer, 50 mM NaCl, 5 mM DTT, 0.1% PEG 6000, pH 6.0 and 50 mM acetic acid buffer, 50 mM NaCl, 5 mM DTT, 0.1% PEG 6000 adjusted to pH 4.0. The inhibitor concentration span was optimized in initial screening experiments for each inhibitor and assay buffer. Reaction data were fitted to the one phase association formula in GraphPad Prism 9 software: Y=Y_0_ + (Plateau-Y_0_)*(1-exp(-K*x)) where in this case Y_0_ and Y represent fluorescence signal at times 0 and *t*, respectively. K represents the observed reaction rate k_obs_ and x the inhibitor concentration. Inactivation rates at each inhibitor concentration were obtained by subtracting the inactivation observed in the control sample: k_obs_ - k_ctrl_. Measurements were performed in three or four independent duplicates or two independent quadruplicates.

## 3. Results

### 3.1. Quantification of anti-SARS-CoV-2 activity in Vero E6 cells

Prior to *in vitro* and structural investigation of the compound interactions with activated CatL, an anti-viral assay was set up in Vero E6 cells. Analyzing the anti-SARS-CoV-2 activity of the selected compounds based on GFP-fluorescence indicated IC_50_ values ranging from the low µM to the low nM regime after 24 and 48 h of incubation (Figure 1A). CI-XII being the only ketoamide in the set of compounds displayed the highest anti-viral potency and the lowest average IC_50_ value after 48 h, i.e. 146 nM (Figure 1A).

**Figure 1.**
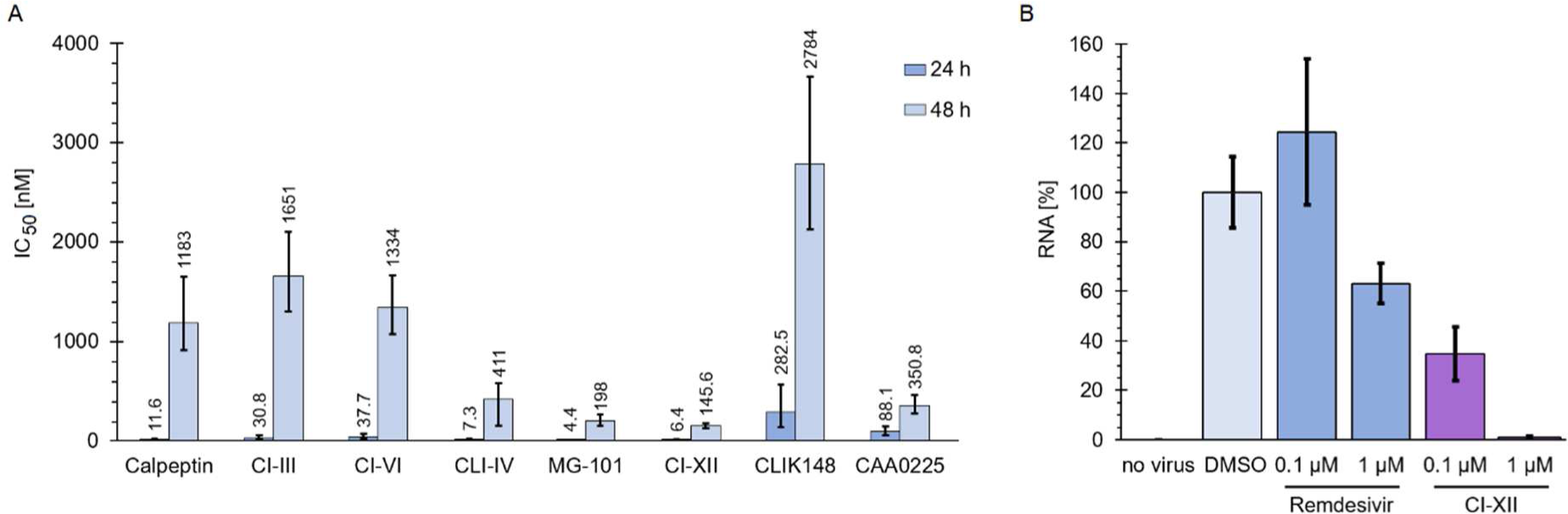
(**A**) CatL inhibitors counteract viral replication of SARS-CoV-2. Vero E6 cells were pretreated with a dilution series of different CatL inhibitors for 1 h and inoculated with SARS-CoV-2-GFP at an MOI of 0.05. IC_50_ values were calculated from live cell imaging data collected using an EssenBioscience Incucyte S3 at 24 h and 48 h post inoculation. Data of four independent experiments with biological triplicates are shown, i.e. n = 4 and m = 3. The boarders of a 95% confidence interval are shown. (**B**) Impact of CI-XII on virus replication was validated using qRT-PCR. Remdesivir was used as a treatment control. Vero E6 cells were inoculated with SARS-CoV-2 Omicron variant BA.1 at an MOI of 0.05. Shown are relative RNA levels relative to the compound-free DMSO control and the respective standard deviation. Depicted are data determined with biological triplicates.

In comparison to CI-XII, the two aldehydes MG-101 and CatL inhibitor IV have similarly low IC_50_ values. These three compounds possess an IC_50_ value below 10 nM after 24 h of cell incubation and below 500 nM after 48 h. The latter is also true for the epoxide CAA0225. Calpeptin has similar IC_50_ values in comparison to CI-VI and stronger inhibition of replication than the rather weakly inhibiting epoxide CLIK148 (Figure 1A).

Due to the high inhibitory potency in the SARS-CoV-2-GFP inhibition assays, CI-XII was further tested against the Omicron variant BA.1 of SARS-CoV-2 in a qRT-PCR experiment (Figure 1B) allowing to quantify viral RNA. The quantity of RNA was reduced by more than 50% in the presence of 100 nM of CI-XII after 24 h. Hence, this approach verifies an IC_50_ value below 100 nM.

### 3.2. Comparison of CatL inhibitor complexes using nanoDSF

To study the stability of protein ligand complexes, i.e. the affinity of ligands, by thermal stability, we used nanoDSF-based screening of the inflection points of denaturation (T_m_) for the CatL inhibitor complexes as shown in Figure 2. CatL was recombinantly produced in *Pichia pastoris*. For pure monomeric CatL (Figure S2), T_m_ was determined to 363.3 ± 0.1 K, buffered at a nearly physiological lysosomal pH value of 5.0. Addition of a 20-fold molar amount of aldehyde and ketoamide compounds resulted in an increase of T_m_ by ∼18 K, whereas the epoxide compounds increased T_m_ by 12-15 K (Figure 2).

**Figure 2.**
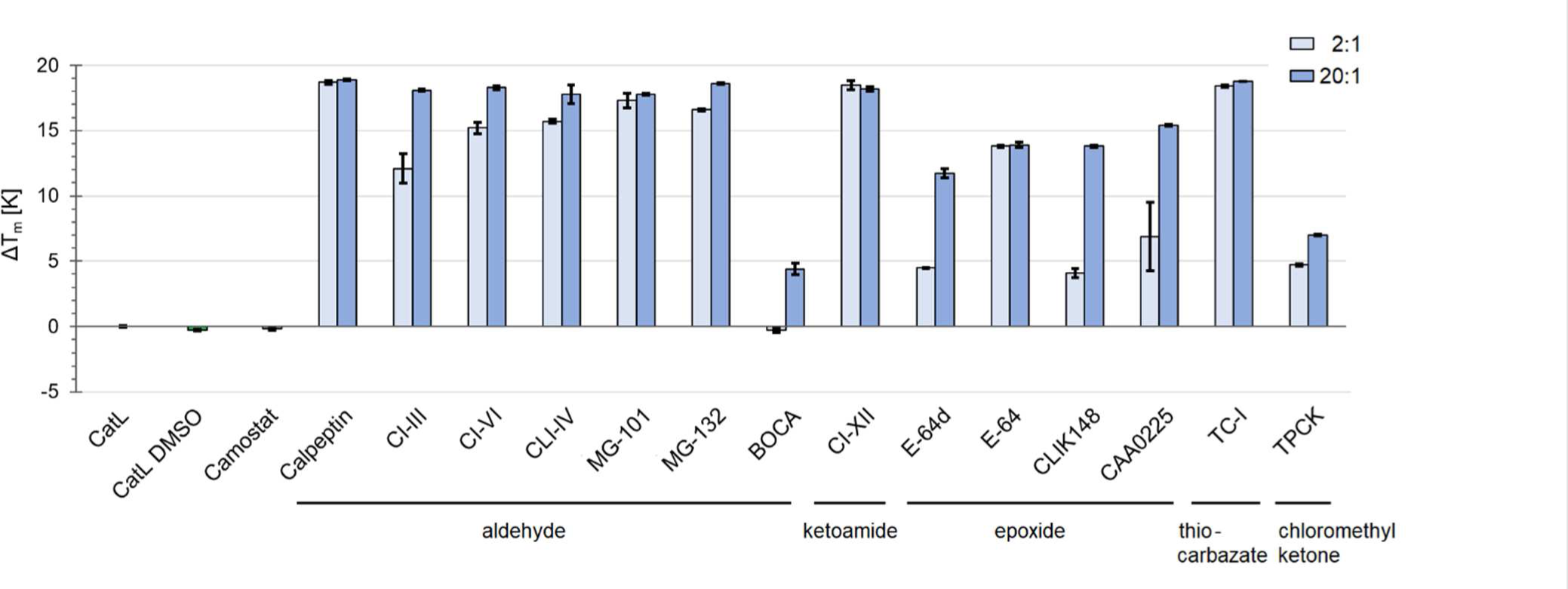
NanoDSF assay. Comparison of thermal stability of CatL as a relative measure for compound affinity. 2:1 and 20:1 mixing ratio of compound to protein are shown. Values for apo protein in buffer without and with 2% DMSO are shown for comparison and as reference for the melting temperature differences (ΔT_m_). Camostat (20:1) as a serine protease inhibitor, was included as an additional negative control.

At a reduced compound to protein mixing ratio of 2:1, the highest affinity to CatL is indicated for the ketoamide CI-XII and the aldehydes calpeptin and MG-101. Those three compounds are, notably, also interacting with the SARS-CoV-2 M^pro^ (Figure S3). TC-I is in the same ΔT_m_ range. The affinity of the tested epoxides to CatL, particularly for E-64 and E-64d is weaker in comparison to CI-XII, indicated by a ΔT_m_ gain reduced by more than half for a twofold molar amount of the respective compound. Overall, nanoDSF data indicated interaction of CatL with all tested compounds (Figure 2), except for the negative control serine protease inhibitor camostat. Consequently, all these compounds were used to set up crystallization experiments. Complementary IC_50_ and K_i_ value references for *in vitro* CatL inhibition are compiled in table S1.

### 3.3. Structural investigation of the compound binding sites

#### 3.3.1. CatL crystal structure

The crystal structures of CatL were determined at maximum resolutions ranging from 1.4 to 1.9 Å. The asymmetric unit (ASU) contains four protein molecules arranged as a distorted tetrahedron without remarkable contact surface areas, which agrees with the observed monomeric state in solution (Figure S2). Beside conformational differences of a loop ranging from amino acid residues 174 to 180 (Figure S4), the four molecules are well superimposable with RMSD over C_alpha_ atoms below 0.4 Å. Induced fit upon inhibitor binding is not observed, because superimposition of all inhibitor complexes with native CatL showed no conformational changes of active site amino acid side chains. In proximity to the S2’ and S3 subsites, electron density maps occasionally showed PEG molecules of varying length, some are reminiscent of crown ethers, interacting with CatL via hydrogen bonds and hydrophobic interactions.

#### 3.3.2. Aldehyde and ketoamide inhibitors

The covalently bound aldehyde BOCA comprises a minimalistic “core-fragment” interacting with the S1 and S2 subsite of CatL related to the scaffold of bigger peptidomimetic aldehyde inhibitors (Figure S5). The aldehyde-type inhibitors, i.e. BOCA, CI-III, CI-VI, CLI-IV, MG-101 and MG-132 as well as the α-ketoamides CI-XII and 13b are covalently bound to the active site Cys25 of CatL forming a thio-hemiacetal or – in case of the ketoamides – a thio-hemiketal. The resulting stereo center of the complexes with the newly formed hydroxyl group attached (see also figure S1 A, B) appeared in the R-configuration. As shown in figures 3 and 4, this enabled the thio-hemiacetals to form an additional hydrogen bond with the side chain amide nitrogen of Gln19 with the interatomic distances ranging from 2.6 to 3.2 Å and 3.3 Å in case of CLI-IV. In the thio-hemiketal of CI-XII and 13b the new hydroxyl group points away from Gln19, but a hydrogen bond to the imidazole of His163 appears (2.7 Å and 2.6 Å respectively; Figure 5). The hydrogen bond to Gln19 is retained by the carbonyl group next to the chiral center (2.6 Å). Figure 3 provides a view of the binding sites of the aldehydes CI-III, VI and CLI-IV. In Figures 4 and 5 the binding sites of the remaining carbonyl-type compounds are shown.

**Figure 3.**
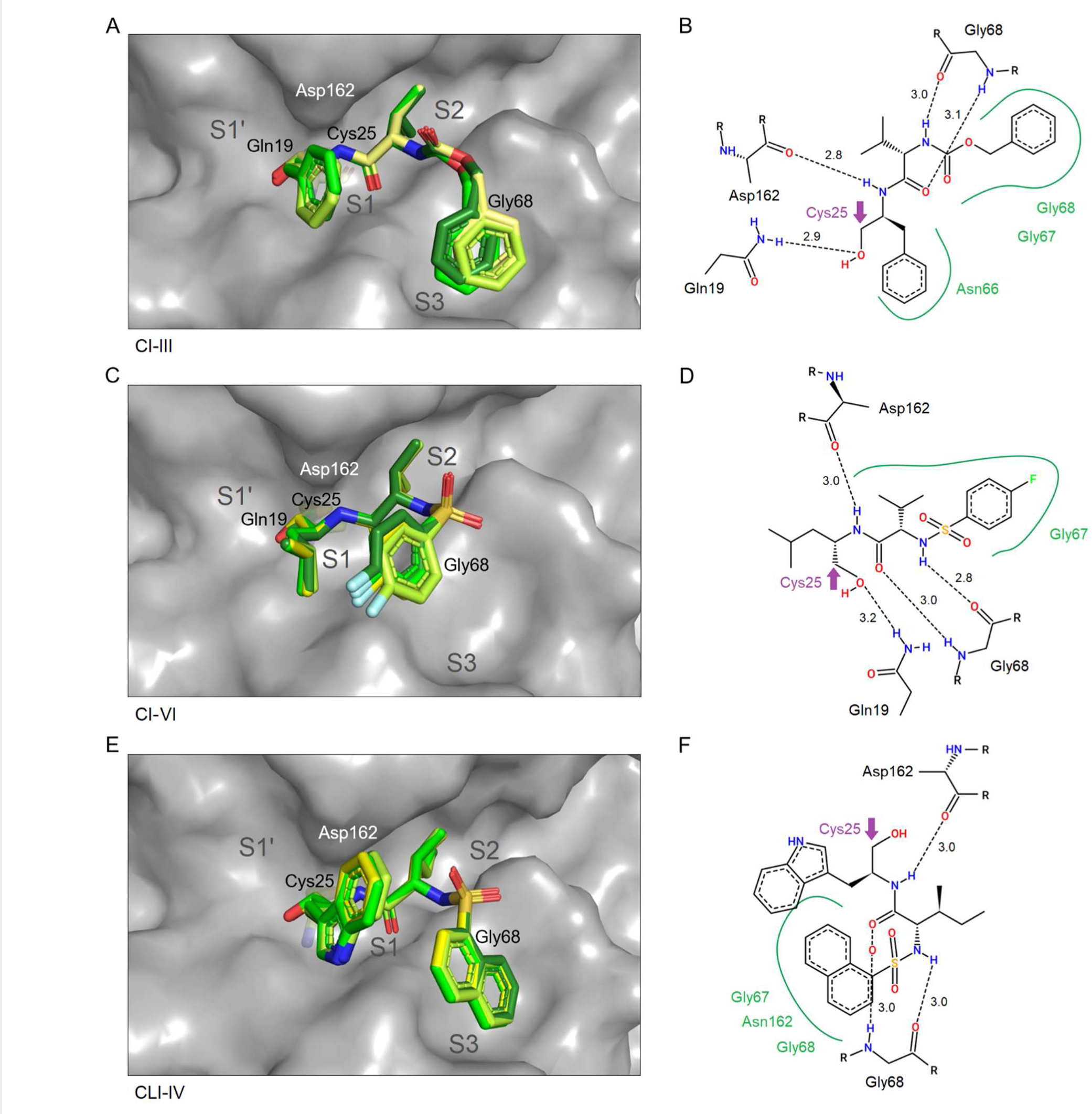
Binding site illustration of *CI-III* (**A, B**), *CI-VI* (**C, D**) and *CLI-IV* (**E, F**). Compound positions for all four molecules in the ASU were superposed for comparison in the panels on the left side (**A, C, E**). The compound molecules are colored from dark green (chain A) fading to yellow (chain D). CatL (chain A) is shown with grey surface representation and Cys25 is indicated using stick representation. Two-dimensional schematic plots of the compound interaction are according to Poseview (**B, D, F**). A purple arrow denotes the thio-hemiacetal with the active site Cys25 of CatL.

**Figure 4.**
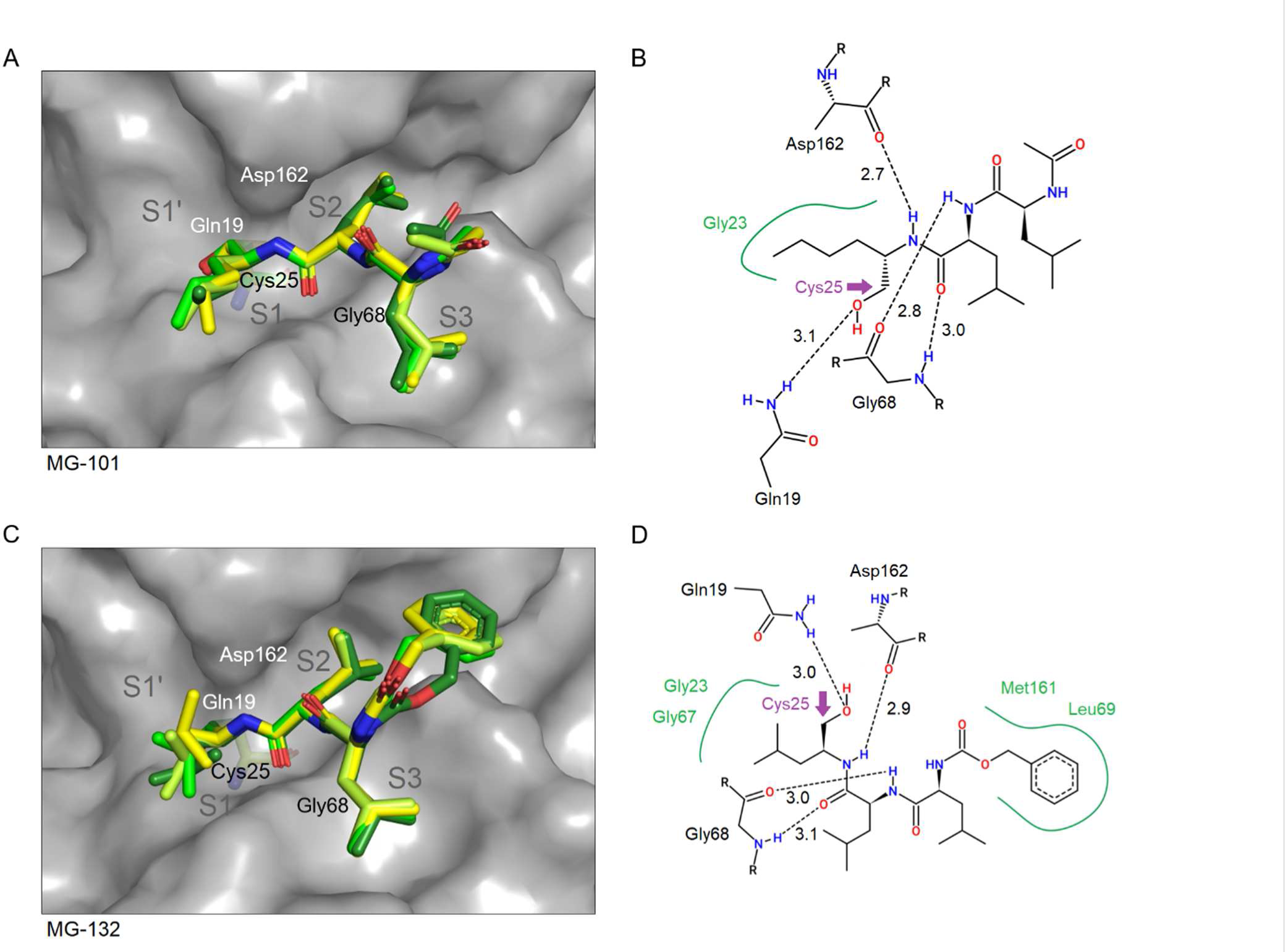
Binding sites of the peptidomimetic aldehydes *MG-101* (**A**, **B**) and *MG-132* (**C**, **D**). Compound positions for all four molecules per ASU were superposed for comparison in the panels on the left side. CatL (chain A) is shown with grey surface representation and Cys25 is shown as sticks. Two-dimensional schematic plots of the compound interaction sites are according to Poseview (**B**, **D**) and a purple arrow denotes the covalent link with the active site Cys25.

**Figure 5.**
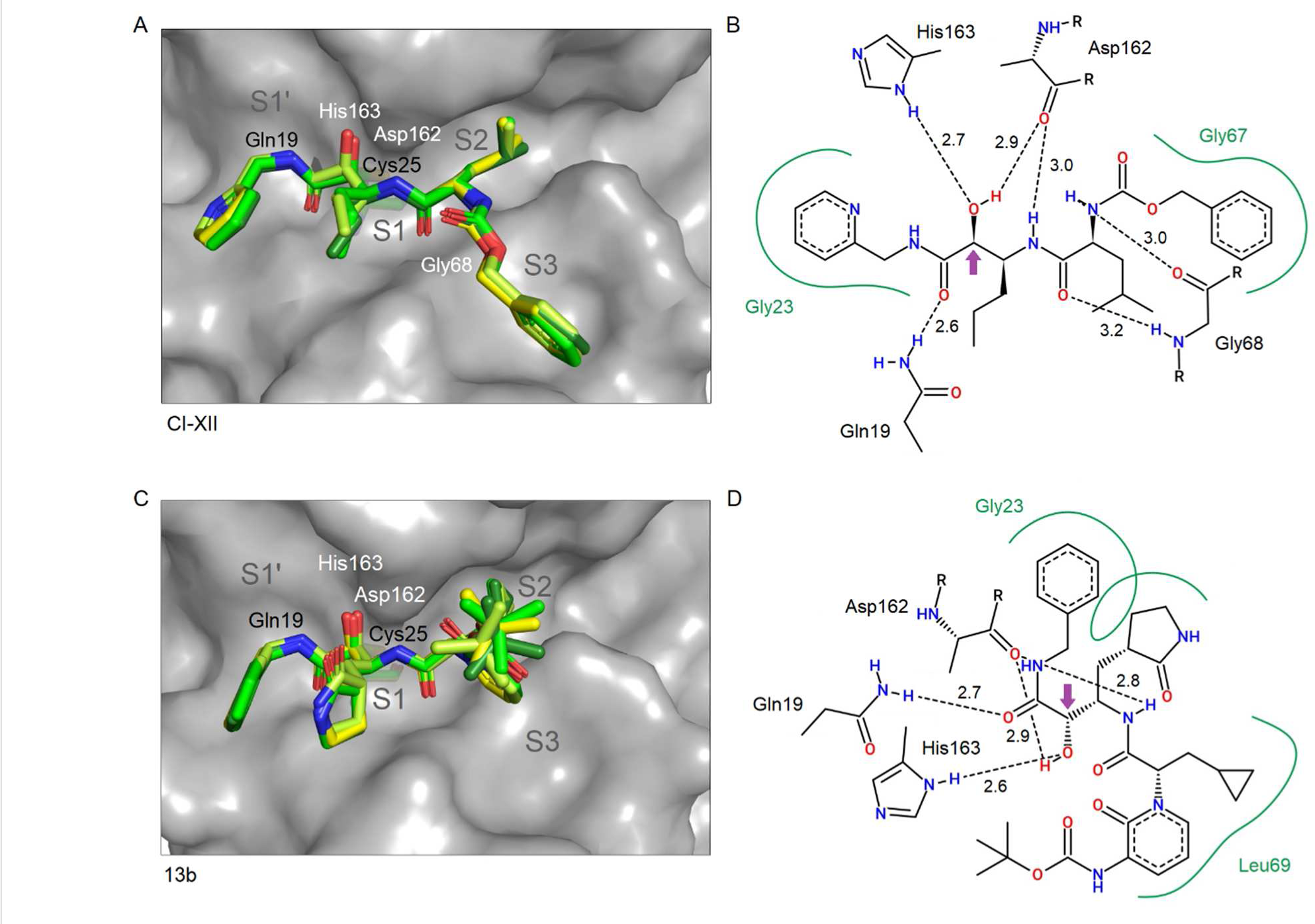
Binding site illustration of the α-ketoamides *CI-XII* (**A**, **B**) and *13b* (**C**, **D**). Compound positions for all four molecules per ASU were superposed for comparison in the panels on the left side. CatL (chain A) is shown with grey surface representation and Cys25 is indicated using stick representation. Two-dimensional schematic plots of the compound interaction sites are according to Poseview (**B**, **D**) and a purple arrow denotes the covalent link with the active site Cys25.

#### 3.3.3. Epoxide-type inhibitors

Both chiral centers of the epoxide rings of the used compounds are in the S-configuration. In the crystal structures, covalent binding of the succinyl epoxide with cathepsin L was observed for E-64, E-64d and CLIK148, whereas non-covalent binding was observed for CAA0225. Hence, to compare and verify the inactivation of CatL by CAA0225 and the structurally related CLIK148 and E-64, a separate enzyme assay was performed (Figure S6). It revealed that CAA0225 is the most potent of the three epoxides, followed by E-64 and CLIK148. Epoxide ring opening and subsequent thioether formation is a result of the nucleophilic addition of Cys25 to an epoxide carbon. The former epoxide oxygen is converted to a hydroxyl substituent and – distinct from the hemiacetals – remains solvent exposed in the covalent complex.

Both carbonyl groups of the succinyl epoxide moiety of E-64, E-64d, CAA0225 and CLIK148 are hydrogen bond acceptors for the amides of the Gln19 side chain and the Gly68 main chain. The terminal carboxylate of E-64 forms a salt bridge with the imidazole of His163. The S2 subsite is bound by hydrophobic side chains, i.e. Leu of E-64d and E-64 or Phe of CLIK148 and CAA0225. The S3 subsite for E-64d and CAA0225 interacts with the terminal phenyl- and iso-pentyl group, respectively, whereas the terminal guanidinium group of E-64 is solvent exposed and faces the S3 subsite with its n-butyl linker (Figure 6).

**Figure 6.**
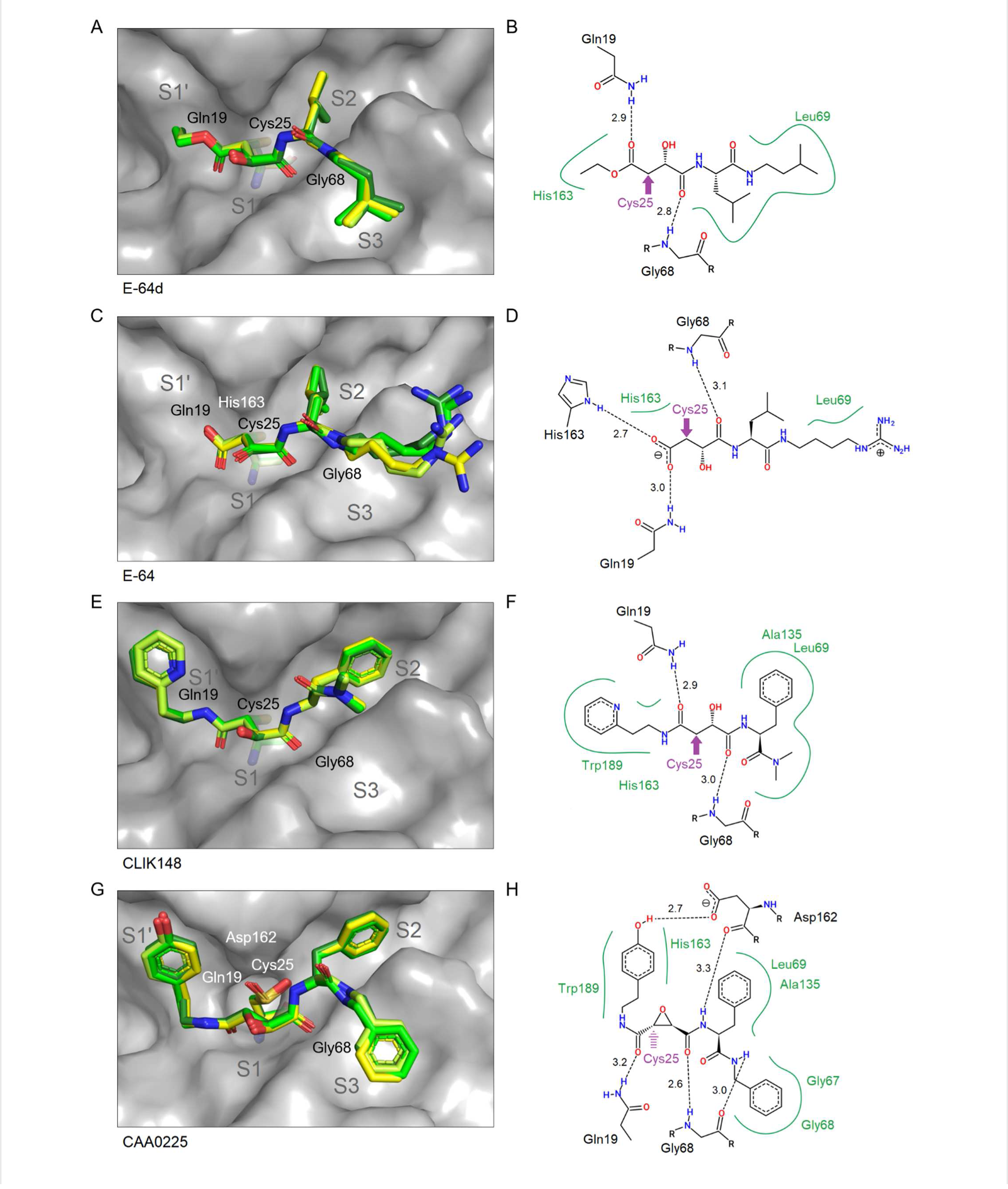
Binding site illustration of the succinyl-epoxides *E-64d* (**A**, **B**), *E-64* (**C**, **D**), *CLIK148* (**E**, **F**) and *CAA0225* (**G, H**). Compound molecules of all four complexes in the ASU were superposed in the panels on the left side (**A, C, E, G**). CatL (chain A) is shown with grey surface representation and Cys25 is shown with stick representation. Two-dimensional schematic plots of the compound interaction with CatL (chain A) are according to Poseview (**B, D, F, H**). A purple arrow denotes the indicated Cys25 linking position upon binding.

At the S1’ subsite hydrophobic interactions of the phenolic moiety of CAA0225 and similarly the pyridine ring of CLIK148 with Trp189 and also His163 are observed. However, the phenolic hydroxyl group of CAA0225 contributes an additional 2.7 Å hydrogen bond with the carboxylate of Asp162, which is unique among the investigated compounds (Figure 6). E-64 and E-64d do not interact with the S1’ subsite or a rather hydrophobic area around Leu69.

#### 3.3.4. A thiocarbazate and a chloromethyl ketone inhibitor

The structures of CatL in complex with the thiocarbazate TC-I and the chloromethyl ketone TPCK are shown in Figure 7. For TC-I the nucleophilic attack of the Cys25 thiolate on the carbonyl carbon next to the hydrazine group induces a substitution reaction resulting in the observed covalent complex with CatL in all four CatL protomers of the crystal and replacing the sulfur containing part of the inhibitor. As a result, the thiol fragment containing the 2-ethylanilino-2-oxoethyl group was not observed in the electron density maps and, thus, characterized to be the leaving group.

**Figure 7.**
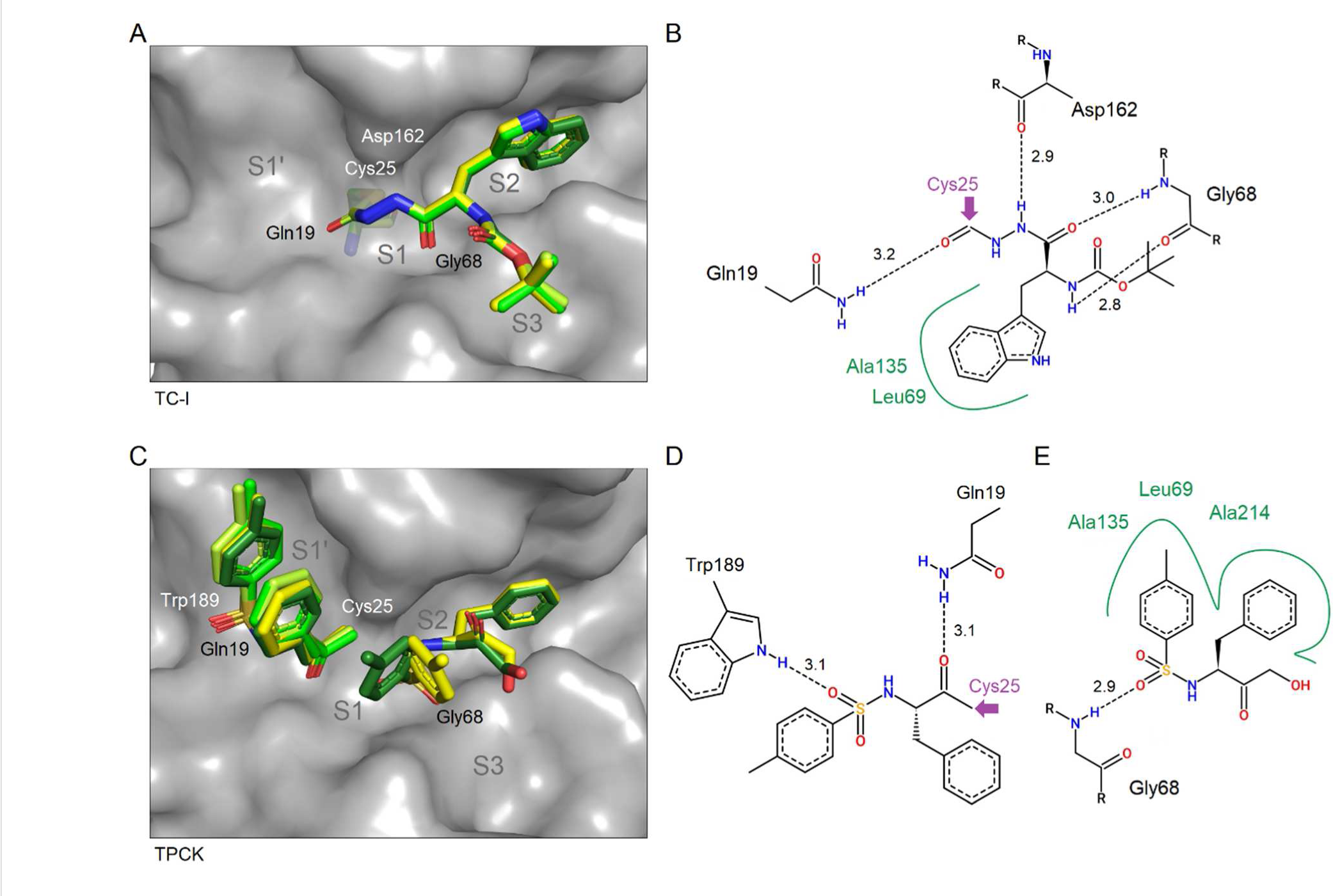
Binding sites and superimposition of *TC-I* (**A**) and *TPCK* (**C**) from the four individual CatL molecules in the ASU. Two-dimensional interaction plots of TC-I (**B**), TPCK covalently bound to the active site (**D**) and a neighboring non-covalently bound molecule in close proximity (**E**) are shown according to chain A. A purple arrow denotes the covalent link with the active site Cys25. Note the stacking of three aromatic rings along subsites S1’ to S1 and the distinct interaction of covalently bound TPCK with the substrate’s C-terminus binding site.

TPCK with its chloromethyl ketone warhead is covalently bound to the active site cysteine forming a thioether linkage. The conformation of the bound inhibitor shows intramolecular aromatic stacking of its tosyl- and phenyl rings. This stacking can be extended by a tosyl moiety of a second TPCK, which binds non-covalently to the S1 and S2 subsites in chains A and D of the ASU (Figure 7C-E). The non-covalently binding TPCK is stabilized by a hydrogen bond with Gly68 and was modelled as a hydrolysis product, due to the missing electron density of the chlorine atom of the warhead.

## 4. Discussion

### 4.1. Quantification of anti-SARS-CoV-2 activity in Vero E6 cells

Most of the compounds under investigation, although to different extends, reduced the propagation of SARS-CoV-2 in Vero cells. The low nanomolar IC_50_ values determined for CI-XII agree with the nanomolar IC_50_ value reported for CI-XII after 3 days of incubation by a viral yield reduction assay using another SARS-CoV-2 strain as well as another detection principle.^47^

Although for example the peptidomimetic aldehyde MG-132 was used for subsequent X-ray crystallography experiments, it was excluded from the *in cellulo* experiments due to notably high toxicity. Toxicity in this case is likely originating from 26S proteasome inhibition with K_i_ in the nanomolar range^48^ and in general can originate from other off-target effects. However, among other fields of application MG-132 was suggested as a suitable tool to selectively analyze the ubiquitin-proteasome pathway in intact cells^49^ and there is some structural similarity to the aldehyde MG-101, which indicated distinct anti-SARS-CoV-2 activity in the same range as CI-XII (Figure 1A). MG-101 has been considered as a drug to support colon cancer prevention^50^ and the comparably small hydrophobic leucine sidechains may contribute to beneficial inhibition of a number of related cysteine proteases in this context.

### 4.2. Two-step mechanism of inhibitor binding

Unexpectedly, in contrast to the other compounds, a non-covalent interaction state of the epoxide CAA0225 with CatL (Figure 6G, H) was unambiguously observed in the electron density maps. The Cys25 sulfur, oxidized to a sulfinic acid, is approximately equally distant to both CAA0225 epoxide ring carbon atoms (3.2 and 3.4 Å respectively, averaged over chains A-D), which are the supposed target for nucleophilic attack and subsequent covalent binding and inactivation as confirmed by the *in vitro* inactivation assay (Figure S6). This suggests that we notably observed the non-covalent intermediate, which is formed in the first step of the reaction. Namely, the covalent inhibitor complexes are expected to form in a two-step mechanism, where the first is reversible and non-covalent in agreement with the following reaction scheme (E: enzyme, I: inhibitor).

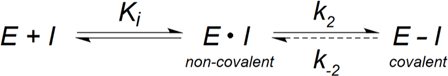

The covalent binding in the second step of the reaction pathway is irreversible for epoxides, thiocarbazates and halomethyl ketones (indicated by the dashed arrow).^25,51^

We verified the ability of CAA0225 to form a covalent (irreversibly) inactivating Cys25 adduct as observed in the crystal structures of the other succinyl-epoxides in solution according to an enzyme assay. CLIK148 has a comparable molecular structure as CAA0225, but is lacking the phenyl ring interacting with the S3 subsite. The lack of S3 interaction of CLIK148 can likely also explain the reduced affinity to CatL (Figure 2). We consequently compared the inhibition potency of CAA0225 to CLIK148 and E-64. In the crystal soaking experiment, the CatL thioether formation by CAA0225 was obviously hindered and slower than the oxidation of Cys25. This is in line with partial oxidation (∼50%) of the active site cysteine to sulfenic acid observed in all protomers of the native CatL structure (PDB 7Z3T). Considering an acidic pH value of the lysosomal environment, inhibition of CatL by CAA0225 and CLIK148 is stronger at pH 6 than at pH 4 (Figure S6). This is in contrast to E-64, which is a stronger inhibitor at the lower pH of 4. Assuming a pK_a_ value around 4 for the carboxylate group of E-64, a larger fraction is deprotonated in an environment at pH 6. Thus, the higher negative charge of this substituent at pH 6 weakens the electrophilicity of the epoxide carbon and hence reducing the affinity to the reactive cysteine. The observed efficient inhibition of CatL by CAA0225, relative to CLIK148, is supposed to contribute to the lower anti-SARS-CoV-2 IC_50_ value under cellular conditions (Figure 1A).

### 4.3. CatL inhibitor complexes: affinity and inhibition

Looking at the affinity, compounds with a relatively high affinity to CatL in the nanoDSF assay possess a high anti-SARS-CoV-2 activity (Figures 1A and 2). The similar behavior of a few of the peptidomimetic aldehydes, CI-XII and epoxides suggests that CatL is indeed a predominant target of the compounds *in vivo.* A few of them, including CI-XII exhibited additional SARS-CoV-2 M^pro^ inhibition potency, as indicated by nanoDSF (Figure S3). Using complementary activity assays the inhibition of M^pro^ by CI-XII, calpeptin, MG-101 and additional related compounds like CI-II and MG-115 has been investigated including the determination of IC_50_ values in the high nanomolar range.^30,47^

The lower affinity of E-64 and other succinyl-epoxide inhibitors for CatL, compared to e.g. CI-XII according to nanoDSF (Figure 2), agrees with an approximately three-times higher IC_50_ value for E-64^27^ (Table S1). Beside CatL inhibition, E-64 is known to have a broad specificity for inhibiting other cysteine proteases.^21^ In addition to a different specificity of E-64 and related succinyl-epoxide inhibitors, the IC_50_ values previously reported might be one obvious reason for a much higher anti-viral activity of CI-XII compared to E-64 or CLIK148 in the fluorescence-based assay.

For peptidomimetic aldehyde inhibitors like MG-101, MG-132, calpeptin or the distantly related GC-376, the IC_50_ values for *in vitro* CatL inhibition down to a lower nanomolar range were reported.^27,28,52^ In comparison to CI-XII, both CI-VI and CLI-IV are slightly lower in affinity. Despite the structural differences, the anti-SARS-CoV-2 activity of CI-VI is also in the same range as for CI-XII. This approximately fits with identical IC_50_ values of 1.6 nM determined previously for CI-VI and CI-XII (Table S1), which is also in the same range as the 1.9 nM determined for CLI-IV.

Only minor chemical modification of MG-101, i.e. changing the norleucine moiety to a methionine by replacing the δ-carbon with a sulfur atom, results in the structure of CI-II, which has not been further investigated, but may adopt a highly similar binding pose and affinity in complex with CatL. Sasaki *et al.* determined a K_i_ value of 0.6 nM for CI-II and a similar K_i_ of 0.5 nM for MG-101.^28^ For comparison, this is higher than K_i_ values determined for CatL inhibition by the two distinct aldehydes calpeptin (0.13 nM) and GC-376 (0.26 nM)^51^, but in the same order of magnitude.

### 4.4. Comparative compound binding

Recently, proteomics-based screening of peptidyl substrates of the cysteine cathepsins B, F, K, L, S and V revealed that CatL positions from P1 to P3 and P1’ are specific for substrate binding, preferably hydrophobic residues^16^. The P2 residue points into the protein and has exceptional preference for nonpolar groups, due to the shape of the hydrophobic S2 subsite. The P1 residue has no obvious contact to the protein surface and has additional probability to tolerate polar uncharged groups. The distinct hydrophobicity, especially in the S2 and S3 subsites, is in favor of different related hydrophobic side chains of the compounds.

The moieties positioned in the **S1’ subsite** are diverse, but only CI-XII, 13b, CLIK148, CAA0225 and TPCK are clearly utilizing the S1’ subsite for interaction to gain affinity and potentially also specificity. Shenoy *et al.*^53^ designed and investigated inhibitors with biphenyl side chains to cover the S’ subsites. They compared conformationally mobile substituents, like biphenyl, with the rigid naphthyl group and pointed out that large rigid substituents like naphthalene are disfavored due to increased corresponding entropic costs of cathepsin inhibition and thereby limiting an improvement of the compound potency.^53^ On the other hand larger contact surface of the inhibitor favors the entropic term for inhibitor affinity. Both aspects must be considered in case of further “modular” peptidomimetic enlargement of a CatL inhibitor inspired by interactions observed in the presented crystal structures. Further extension and modifications along P1’ and P2’ are facilitated with the epoxide and ketoamide warheads, but impossible with terminal aldehyde warheads.

The structurally closely related compounds CLIK148 and CAA0225 bind to the S1’ subsite with their pyridine and phenolic ring, respectively, in a very similar orientation. The phenolic hydroxyl group of CAA0225 is hydrogen bond donor to the carboxylate of Asp162 (2.7 Å). Due to its phenolic moiety in S1’ position, CAA0225 is the only compound under investigation forming a hydrogen bond with the side chain of Asp162. This is rendering the option to further optimize lead compounds, which specifically occupy the S1’ or potentially even the S2’ subsite. The tosyl substituent of the covalently bound TPCK is also in the S1’ subsite like CLIK148 and CAA0225. The pyridine moiety of CI-XII (Figure 5 A, B) points into the opposite direction of the aromatic rings discussed above towards the S2’ subsite due to a ∼180° rotation at the linking methylene group. The phenyl ring in P1’ position of 13b is in a similar position as the pyridine ring of CI-XII. The indole moiety of the tryptophan side chain of CLI-IV (Figure 3 E, F) seems to be too large to occupy the S1’ subsite and bins in a solvent exposed orientation.

The ethyl-pyridine moiety of CLIK148 is interacting with the S1’ subsite. Considering some rotational flexibility of the ethyl linker, the pyridine ring might be able to occupy a different position around the S’ subsites overlapping with a PEG molecule from the solvent identified at this position in the CLIK148 complex structure. This position would then be much more similar to the pyridine binding position of CLIK148 when binding to papain.^54^ Binding of CLIK148 to papain is supported by hydrophobic interaction of the pyridine ring with Trp177 and Gly23, but the broad subsite of papain does not provide additional specific interaction at this position. Within CLIK148 the pyridine nitrogen atom is hydrogen bond acceptor (2.9 Å) for the intramolecular amide N-H of its own linker. Nonetheless, there is hydrophobic and weak T-shaped ring stacking interaction with the indole of Trp189 to keep the pyridine in position. On the opposite side of CLIK148, in the S2 subsite, the phenyl moiety is covered by hydrophobic interactions. Potentially, a solvent-exposed hydroxylation in *para*- or *ortho*-position to provide a hydrogen bond donor to carbonyl Met161 could be added to the phenyl ring located in the S2 subsite.

For several of the investigated compounds (Table 1), the **S1 subsite** with the reactive site cysteine is occupied by a small alkyl moiety, whereas the narrow **S2 subsite**, i.e. the major specificity-determining subsite among cathepsin endopeptidases, is in most cases covered by a variety of small aliphatic or aromatic side chains. A dedicated isopropyl side chain, e.g. found in CI-XII, E-64d and E-64, may be increased in size to potentially fit the S2 subsite more efficiently. The distinct cyclopropyl moiety of α-ketoamide 13b binding to and optimized for the S2 subsite of SARS-CoV-2 M^pro^^36^ is located in the S2 subsite of CatL as well (Figure 5C, D). Highly similar to the epoxide CAA0225, a phenyl ring of CLIK148 is bound in the S2 subsite, resembling the native substrate specificity of CatL. The phenyl ring is held in position via hydrophobic interaction with Leu69 and Ala135 (Figure 6E, F). The preference of CatL for aromatic rings at the P2 position is also supported by the phenyl moiety of the non-covalent TPCK observed in the S2 subsite. In the case of TC-I, an even larger aromatic moiety, i.e. an indole, well fits in the S2 subsite (Figure 7A, B).

There is a lot of variation of the **S3 subsite** binding moieties among the inhibitors (Figure 8). The hydrophobic S3 subsite is essentially formed by Leu69 and Tyr72, which correspond to Phe69 and Arg72 respectively in the tissue-specific CatV. The S3 subsite of CatL is typically interacting with a hydrophobic moiety, e.g. the naphthyl ring of CLI-IV or the phenyl ring of CI-III, which is similarly also found in calpeptin. MG-101, MG-132 and E-64d possess a smaller isopropyl-group to bind at the S3 position. For TC-I the corresponding hydrophobic moiety is enlarged to a *tert*-butyl group, which however does not provide additional interaction with CatL compared to the isopropyl-group at this position.

**Figure 8.**
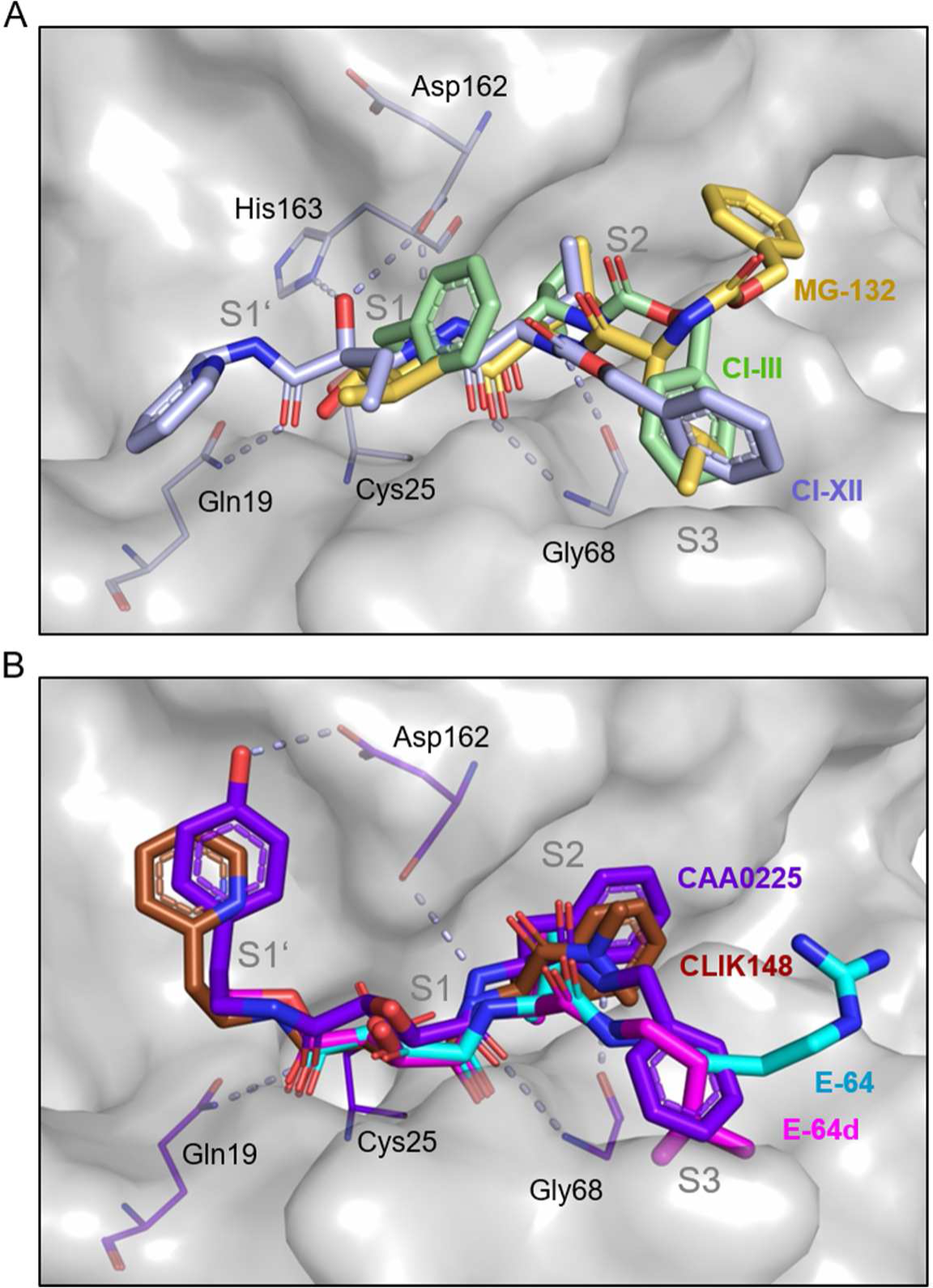
Compound superimposition shows conserved hydrogen bonding with the main chain atoms of amino acids Gln19, Gly68 and Asp162 of CatL and indicates ways to complement individual moieties in different subsites: (**A**) *MG-132* (yellow), *CI-III* (green) and *CI-XII* (light blue) as well as (**B**) the epoxides *CAA0225* (purple), *CLIK148* (brown), *E-64* (cyan) and *E-64d* (magenta) as individually shown in figures 3-6. Compounds are shown in stick representation, surrounding amino acid residues as lines (chain A of the ASU). Dashed lines indicate hydrogen bonds of CI-XII in panel A and CAA0225 in panel B. Notably, the hydrogen bonds of CI-XII with His163 and CAA0225 with the side chain of Asp162, potentially related to enzyme specificity, are both exclusively observed for the respective compound.

To fit the rather narrow S3 subsite the naphthyl moiety of CatL inhibitor IV is interacting with Gly67 and needs to slightly rotate to fit the pocket widthwise with the ring plane tilted over one pocket side, unlike the much smaller phenyl ring of CI-III or CAA0225. The phenyl ring is however not tightly in position, when comparing the four protein chains, especially for CI-III. In contrast to CI-III and CAA0225, the geometry of CI-XII puts the corresponding phenyl ring in a slightly different position of the same hydrophobic site. This results in ∼90° rotation of the ring, stabilized by π-amide stacking interaction with Glu63. Regarding CI-VI, the characteristic fluorinated phenyl moiety is not similarly positioned in the core of the S3 subsite and the highly electronegative fluorine sticks out at the site boarder and interacts with a solvent water molecule. Some inhibitors, e.g. CI-VI, CLIK148 and E-64 do not possess a dedicated moiety to occupy the center of the S3 subsite and also in the complex with 13b the S3 subsite is empty. However, the scaffolds of CLIK148 and E-64 seem to allow adding another alkyl moiety branching off the compound close to the S3 subsite, potentially increasing their rather low affinity and specificity in comparison to other CatL inhibitors even without changing functional groups of the compound. In comparison of the determined binding sites, widely shared hydrogen bond hotspots across the subsites of CatL include Gln19, Asp162 and Gly68 as highlighted individually in figures 3-7 and S5 as well as figure 8.

### 4.5. Specificity of cathepsin L inhibitors

Inhibition of related cysteine-type cathepsins by the investigated compounds is primarily explained by a high level of sequence and structural similarity, including their active sites. This is generally a major limitation of the therapeutic potency of inhibitors targeting mammal proteases. Most of the investigated compounds were not optimized for specific inhibition of CatL. In comparison to human CatL, the cathepsins CatS (PDB 1GLO), CatK (PDB 5TUN), CatB (PDB 2IPP) and the highly tissue-specific CatV (PDB 7Q8I/7Q8O/7Q8Q) possess overall RMSD values (Cα) below 1.2 Å. Both L-domain loops (amino acid residues 19-25 and 61-69) involved in substrate binding are however only partly conserved. For example, the mutation of Leu69 in CatL to a tyrosine in CatB and CatK and to phenylalanine in CatV, CatS and other cathepsins could be utilized to gain specificity for the investigated compounds. Further the exchange of Tyr72 to the corresponding Arg72 in CatV is relevant for the shape of the S3 subsite. A detailed sequence alignment and comparison of substrate specificity was provided by Turk and Gunčar.^55^ Due to the involvement of different related cysteine proteases in SARS-CoV-2 infected cells and being aware of potential off-target effects, anti-viral drugs may however benefit from inhibiting multiple closely related cathepsins.

In combination with host-cell proteases like cathepsins, the coronaviral protease M^pro^ has been suggested for dual targeting due to structural similarity. For example, the pyridine ring of CI-XII is interacting with the S1’ subsite of CatL mainly via Gly23 (Figure 5A, B). The nitrogen atom of this pyridine ring is not involved in specific interaction with CatL, but in case of M^pro^ it forms a hydrogen bond with the sidechain of SARS-CoV-2 M^pro^ His163^33^ in proximity to the catalytic His41 and is thereby contributing to a multi-targeting concept of the compound.

The covalent thio-hemiketal formed upon binding of CI-XII to the active site cysteine of CatL adopts an R-configuration as also observed for 13b. An R-configuration is also observed for CI-XII binding to SARS-CoV-2 M^pro^, which is distinct from other α-ketoamide inhibitors, including 13b, which bind to SARS-CoV-2 M^pro^ with S-configuration.^33,36^ The tripeptide sulfone inhibitor GC-376, in part structurally similar to MG-132 and other peptidomimetic inhibitors, actually adopts either the S- or R-configuration at the active site of SARS-CoV-2 M^pro^ in different protomers within the same X-ray crystal structure.^33^ The CatL crystal structures presented here show however no indication for alternative configuration among the four individual protomers of the ASU, suggesting a more specific active site recognition.

### 4.6. Drug development outlook

We have determined the crystal structures of 14 compounds in complex with CatL at high resolution. Ten of these compounds have additionally been tested for anti-viral activity against SARS-CoV-2. Seven compounds, including four aldehydes, two epoxides, and one ketoamide, indeed reduced viral replication in Vero-E6 cells.

Due to their adverse pharmacokinetic properties and toxicity side effects, aldehydes are regarded as unfavorable for drug development.^56^ Nevertheless, rather simple chemical modifications of this aldehyde warhead, i.e. a sulfonic acid moiety applied to the compound GC-376 or self-masked aldehyde inhibitors^57^ have allowed to widely circumvent this problem and could be applied to the inhibitors from the present work. Compounds with alternative warheads, such as the keto-amide CI-XII possess comparably high anti-SARS-CoV-2 potency in Vero E6 cells, and qualify for further evaluation in mouse or Siberian hamster models. CI-XII already indicated an acceptable cytotoxicity (CC_50_ > 50 µM) and a half maximal effective concentration below 1 µM in another assay using Vero cells and the SARS-CoV-2 wild type SA-WA1/2020.^47^ In the context of dual-targeting, the keto-amide CI-XII and related CatL inhibitors showed off-target inhibition of viral cysteine proteases like M^pro^.^27,58^ Inversely, inhibitor 13b, which was developed as a highly specific M^pro^ inhibitor, also binds to CatL.

Further antiviral activity of the investigated calpain inhibitors in viral infections might occur from the inhibition of calpain itself.^34,59^ It has been postulated that calpain inhibition interferes with the clathrin coat formation for the vesicles required for endosomal cell entry of the virus. In combination with cathepsin inhibition, this effect would then hinder endosomal cell entry even stronger and should be investigated for its potential to reduce pulmonary fibrosis originating from a SARS-CoV-2 infection.^31^

In addition, some of the CatL inhibitors have been proposed for therapeutic potential in other contexts. E-64d, derived from the *Aspergillus japonicus* secondary metabolite E-64, showed some pharmaceutical potential to treat Moloney murine leukemia virus.^60^ An immune response potentiation in *Leishmania major* infections for CLIK148^61^, anti-parasite properties of a CatL thiocarbazate inhibitor^62^ and improvement of cardiac function in reperfusion injury by CAA0225^63^ have been reported.

While viral proteases, such as SARS-CoV-2 M^pro^, were in focus of recent drug development and screening efforts, host cell proteases, including CatL, present equally potent drug targets. Although additional side effects could occur when targeting host cell enzymes, this approach is less prone to adaptation of the virus over time via mutations that reduce the drug potency. Associated optimization of cathepsin inhibitors includes utilizing the S1’subsite of CatL to gain affinity, utilizing interaction with the few specificity determining residues of CatL and addition of optimized hydrophobic moieties binding in the S2 and S3 subsite. For example, the two neighboring TPCK binding sites may inspire the design of a preliminary TPCK derivative, which is not only interacting with the S1’ subsite of CatL, but extended towards the S2 subsite to cover a larger area of the active site. For CI-XII, low toxicity and the low IC_50_ values determined are already encouraging for future medicinal applications. The available structural data of CI-XII would also allow to “re-balance” dual-targeting of CatL and the coronaviral main protease. Additional investigation of CatL inhibitors *in vitro* and *in vivo* will not only contribute to optimization of an anti-coronaviral drug, but also increase the level of preparedness to deal with cathepsin-dependent viral infections and potentially other diseases of high relevance in the future.

## Acknowledgement

Parts of this research were supported by facilities and Maxwell computational resources provided by Deutsches Elektronen-Synchrotron (Helmholtz-Gemeinschaft; Hamburg, Germany). The authors acknowledge support in diffraction data collection and beamtime coordination by Janina Sprenger and Viviane Kremling (Center for Free-Electron Laser Science, DESY). The authors further acknowledge support by the beamline scientists Johanna Hakanpää, Helena Taberman, Guillaume Pompidor and Spyros Chatziefthymiou at PETRA III beamline P11, where parts of this research were performed as well as support by Stephan Niebling and the EMBL Hamburg sample preparation and characterization facility headed by Maria Garcia Alai.

## Data availability

X-ray crystal structures related to this research will be available in the protein data bank via entry IDs 7QKD, 7ZS7, 7ZVF, 7ZXA, 8A4U, 8A4V, 8A4W, 8A4X, 8A5B, 8AHV, 8B4F, 8C77, 8OFA and 8PRX. Cathepsin L in complex with covalently bound CA-074 methyl ester is available via ID 8OZA.

## Author contributions

Designed research: SF, DT, VT, HNC, SG, GE, AM

Wrote and discussed manuscript: SF, AH, WH, JLo, JLi, WE, DT, AM, with input from all authors

Sample preparation: SF, JLo, KK, AU, NL, AS, AH

Cell infection and replication assays: AH

X-ray data collection and analysis: SF, JLi, PYAR, WH, SG, WE

Performed and analyzed *in vitro* assays: JLo, SF

Provided resources/material: DT, GE, HT, HNC, AM

## References

1. Zhao, M.-M. et al. Cathepsin L plays a key role in SARS-CoV-2 infection in humans and humanized mice and is a promising target for new drug development. Signal Transduct. Target. Ther. 6, 134 (2021).

2. Koch, J. et al. TMPRSS2 expression dictates the entry route used by SARS-CoV-2 to infect host cells. EMBO J. 40, (2021).

3. Hoffmann, M. et al. SARS-CoV-2 Cell Entry Depends on ACE2 and TMPRSS2 and Is Blocked by a Clinically Proven Protease Inhibitor. Cell 181, 271–280.e8 (2020).

4. Zhao, M.-M. et al. Novel cleavage sites identified in SARS-CoV-2 spike protein reveal mechanism for cathepsin L-facilitated viral infection and treatment strategies. Cell Discov. 8, 53 (2022).

5. Padmanabhan, P. & Dixit, N. M. Modelling how the altered usage of cell entry pathways by the SARS-CoV-2 Omicron variant may affect the efficacy and synergy of TMPRSS2 and Cathepsin B/L inhibitors. http://biorxiv.org/lookup/doi/10.1101/2022.01.13.476267 (2022), doi:10.1101/2022.01.13.476267.

6. Liu, T., Luo, S., Libby, P. & Shi, G.-P. Cathepsin L-selective inhibitors: A potentially promising treatment for COVID-19 patients. Pharmacol. Ther. 213, 107587 (2020).

7. Pager, C. T. & Dutch, R. E. Cathepsin L Is Involved in Proteolytic Processing of the Hendra Virus Fusion Protein. J. Virol. 79, 12714–12720 (2005).

8. Pager, C. T., Craft, W. W., Patch, J. & Dutch, R. E. A mature and fusogenic form of the Nipah virus fusion protein requires proteolytic processing by cathepsin L. Virology 346, 251–257 (2006).

9. Islam, M. I. et al. Regulatory role of cathepsin L in induction of nuclear laminopathy in Alzheimer’s disease. Aging Cell 21, (2022).

10. Salpeter, S. J. et al. A novel cysteine cathepsin inhibitor yields macrophage cell death and mammary tumor regression. Oncogene 34, 6066–6078 (2015).

11. Zwicker, J. D. et al. Optimization of dipeptidic inhibitors of cathepsin L for improved *Toxoplasma gondii* selectivity and CNS permeability. Bioorg. Med. Chem. Lett. 28, 1972– 1980 (2018).

12. Garsen, M. et al. Cathepsin L is crucial for the development of early experimental diabetic nephropathy. Kidney Int. 90, 1012–1022 (2016).

13. Turk, V. et al. Cysteine cathepsins: From structure, function and regulation to new frontiers. Biochim. Biophys. Acta BBA - Proteins Proteomics 1824, 68–88 (2012).

14. Mason, R. W., Johnson, D. A., Barrett, A. J. & Chapman, H. A. Elastinolytic activity of human cathepsin L. Biochem. J. 233, 925–927 (1986).

15. Biniossek, M. L., Nägler, D. K., Becker-Pauly, C. & Schilling, O. Proteomic Identification of Protease Cleavage Sites Characterizes Prime and Non-prime Specificity of Cysteine Cathepsins B, L, and S. J. Proteome Res. 10, 5363–5373 (2011).

16. Tušar, L. et al. Proteomic data and structure analysis combined reveal interplay of structural rigidity and flexibility on selectivity of cysteine cathepsins. *Commun*. Biol. 6, 450 (2023).

17. Schechter, I. & Berger, A. On the size of the active site in proteases. I. Papain. Biochem. Biophys. Res. Commun. 27, 157–162 (1967).

18. Drag, M. & Salvesen, G. S. Emerging principles in protease-based drug discovery. Nat. Rev. Drug Discov. 9, 690–701 (2010).

19. Zhang, H. et al. Vinyl sulfone-based inhibitors of trypanosomal cysteine protease rhodesain with improved antitrypanosomal activities. Bioorg. Med. Chem. Lett. 30, 127217 (2020).

20. Citarella, A. & Micale, N. Peptidyl Fluoromethyl Ketones and Their Applications in Medicinal Chemistry. Molecules 25, 4031 (2020).

21. Barrett, A. J. et al. L-*trans*-Epoxysuccinyl-leucylamido(4-guanidino)butane (E-64) and its analogues as inhibitors of cysteine proteinases including cathepsins B, H and L. Biochem. J. 201, 189–198 (1982).

22. Thompson, R. C. [19] Peptide aldehydes: Potent inhibitors of serine and cysteine proteases. in Methods in Enzymology vol. 46 220–225 (Elsevier, 1977).

23. Zhang, L. et al. α-Ketoamides as Broad-Spectrum Inhibitors of Coronavirus and Enterovirus Replication: Structure-Based Design, Synthesis, and Activity Assessment. J. Med. Chem. 63, 4562–4578 (2020).

24. Brewitz, L. et al. Alkyne Derivatives of SARS-CoV-2 Main Protease Inhibitors Including Nirmatrelvir Inhibit by Reacting Covalently with the Nucleophilic Cysteine. J. Med. Chem. (2023), doi:10.1021/acs.jmedchem.2c01627

25. Wolf, W. M. et al. Inhibition of proteinase K by methoxysuccinyl-Ala-Ala-Pro-Ala-chloromethyl ketone. An x-ray study at 2.2-A resolution. J. Biol. Chem. 266, 17695–17699 (1991).

26. Yuan, L. et al. A novel cathepsin L inhibitor prevents the progression of idiopathic pulmonary fibrosis. Bioorganic Chem. 94, 103417 (2020).

27. Hu, Y. et al. Boceprevir, Calpain Inhibitors II and XII, and GC-376 Have Broad-Spectrum Antiviral Activity against Coronaviruses. ACS Infect. Dis. 7, 586–597 (2021).

28. Sasaki, T. et al. Inhibitory Effect of di- and Tripeptidyl Aldehydes on Calpains and Cathepsins. J. Enzym. Inhib. 3, 195–201 (1990).

29. Myers, M. C., Shah, P. P., Diamond, S. L., Huryn, D. M. & Smith, A. B. Identification and synthesis of a unique thiocarbazate cathepsin L inhibitor. Bioorg. Med. Chem. Lett. 18, 210–214 (2008).

30. Günther, S. et al. X-ray screening identifies active site and allosteric inhibitors of SARS-CoV-2 main protease. Science eabf7945 (2021), doi:10.1126/science.abf7945

31. Inal, J., Paizuldaeva, A. & Terziu, E. Therapeutic use of calpeptin in COVID-19 infection. Clin. Sci. 136, 1439–1447 (2022).

32. Reinke, P. et al. Calpeptin is a potent cathepsin inhibitor and drug candidate for SARS-CoV-2 infections. (2023), doi:10.21203/rs.3.rs-2450926/v1

33. Sacco, M. D. et al. Structure and inhibition of the SARS-CoV-2 main protease reveal strategy for developing dual inhibitors against M ^pro^ and cathepsin L. Sci. Adv. 6, (2020).

34. Schneider, M. et al. Severe Acute Respiratory Syndrome Coronavirus Replication Is Severely Impaired by MG132 due to Proteasome-Independent Inhibition of M-Calpain. J. Virol. 86, 10112–10122 (2012).

35. Shah, P. P. et al. A Small-Molecule Oxocarbazate Inhibitor of Human Cathepsin L Blocks Severe Acute Respiratory Syndrome and Ebola Pseudotype Virus Infection into Human Embryonic Kidney 293T cells. Mol. Pharmacol. 78, 319–324 (2010).

36. Zhang, L. et al. Crystal structure of SARS-CoV-2 main protease provides a basis for design of improved α-ketoamide inhibitors. Science 368, 409–412 (2020).

37. Stukalov, A. et al. Multilevel proteomics reveals host perturbations by SARS-CoV-2 and SARS-CoV. Nature 594, 246–252 (2021).

38. Thi Nhu Thao, T., et al. Rapid reconstruction of SARS-CoV-2 using a synthetic genomics platform. Nature 582, 561–565 (2020).

39. Miheliĕ, M., Doberšek, A., Gunĕar, G. & Turk, D. Inhibitory Fragment from the p41 Form of Invariant Chain Can Regulate Activity of Cysteine Cathepsins in Antigen Presentation. J. Biol. Chem. 283, 14453–14460 (2008).

40. Burastero, O. et al. eSPC: an online data-analysis platform for molecular biophysics. Acta Crystallogr. Sect. Struct. Biol. 77, 1241–1250 (2021).

41. Kabsch, W. *XDS*. Acta Crystallogr. D Biol. Crystallogr. 66, 125–132 (2010).

42. Winn, M. D. et al. Overview of the CCP4 suite and current developments. Acta Crystallogr. D Biol. Crystallogr. 67, 235–242 (2011).

43. McCoy, A. J. et al. *Phaser* crystallographic software. J. Appl. Crystallogr. 40, 658–674 (2007).

44. Adams, P. D., et al. *PHENIX* : a comprehensive Python-based system for macromolecular structure solution. Acta Crystallogr. D Biol. Crystallogr. 66, 213–221 (2010).

45. Emsley, P., Lohkamp, B., Scott, W. G. & Cowtan, K. Features and development of *Coot*. Acta Crystallogr. D Biol. Crystallogr. 66, 486–501 (2010).

46. Stierand, K., Maass, P. C. & Rarey, M. Molecular complexes at a glance: automated generation of two-dimensional complex diagrams. Bioinformatics 22, 1710–1716 (2006).

47. Ma, C. et al. Boceprevir, GC-376, and calpain inhibitors II, XII inhibit SARS-CoV-2 viral replication by targeting the viral main protease. Cell Res. 30, 678–692 (2020).

48. Lee, D. H. & Goldberg, A. L. Proteasome inhibitors: valuable new tools for cell biologists. Trends Cell Biol. 8, 397–403 (1998).

49. Lee, D. H. & Goldberg, A. L. Selective Inhibitors of the Proteasome-dependent and Vacuolar Pathways of Protein Degradation in Saccharomyces cerevisiae. J. Biol. Chem. 271, 27280–27284 (1996).

50. Li, S. et al. ALLN hinders HCT116 tumor growth through Bax-dependent apoptosis. Biochem. Biophys. Res. Commun. 437, 325–330 (2013).

51. Konuray, A. O., Fernández-Francos, X. & Ramis, X. Analysis of the reaction mechanism of the thiol–epoxy addition initiated by nucleophilic tertiary amines. Polym. Chem. 8, 5934–5947 (2017).

52. Yang, W.-L. et al. Potential drug discovery for COVID-19 treatment targeting Cathepsin L using a deep learning-based strategy. Comput. Struct. Biotechnol. J. 20, 2442–2454 (2022).

53. Shenoy, R. T. et al. A Combined Crystallographic and Molecular Dynamics Study of Cathepsin L Retrobinding Inhibitors. J. Med. Chem. 52, 6335–6346 (2009).

54. Tsuge, H. et al. Inhibition mechanism of cathepsin L-specific inhibitors based on the crystal structure of papain-CLIK148 complex. Biochem. Biophys. Res. Commun. 266, 411–416 (1999).

55. Turk, D. & Gunčar, G. Lysosomal cysteine proteases (cathepsins): promising drug targets. Acta Crystallogr. D Biol. Crystallogr. 59, 203–213 (2003).

56. LoPachin, R. M. & Gavin, T. Molecular Mechanisms of Aldehyde Toxicity: A Chemical Perspective. Chem. Res. Toxicol. 27, 1081–1091 (2014).

57. Li, L. et al. Self-Masked Aldehyde Inhibitors: A Novel Strategy for Inhibiting Cysteine Proteases. J. Med. Chem. 64, 11267–11287 (2021).

58. Costanzi, E. et al. Structural and Biochemical Analysis of the Dual Inhibition of MG-132 against SARS-CoV-2 Main Protease (Mpro/3CLpro) and Human Cathepsin-L. Int. J. Mol. Sci. 22, 11779 (2021).

59. Carragher, N. Calpain Inhibition: A Therapeutic Strategy Targeting Multiple Disease States. Curr. Pharm. Des. 12, 615–638 (2006).

60. Kumar, P., Nachagari, D., Fields, C., Franks, J. & Albritton, L. M. Host Cell Cathepsins Potentiate Moloney Murine Leukemia Virus Infection. J. Virol. 81, 10506–10514 (2007).

61. Zhang, T. et al. Treatment with cathepsin L inhibitor potentiates Th2-type immune response in Leishmania major-infected BALB/c mice. Int. Immunol. 13, 975–982 (2001).

62. Shah, P. P. et al. Kinetic Characterization and Molecular Docking of a Novel, Potent, and Selective Slow-Binding Inhibitor of Human Cathepsin L. Mol. Pharmacol. 74, 34–41 (2008).

63. He, W., McCarroll, D., Elliott, E. B. & Loughrey, C. M. The Cathepsin-L Inhibitor CAA0225 Improves Cardiac Function During Ischaemia-Reperfusion. Biophys. J. 106, 729a (2014).

