## Supporting material for "Structural elucidation and antiviral activity of cathepsin L inhibitors with carbonyl and epoxide warheads"

### Supplementary material

**Authors**

Sven Falke[1*], Julia Lieske[1], Alexander Herrmann[2], Jure Loboda[3], Sebastian Günther[1], Patrick YA Reinke[1], Wiebke Ewert[1], Katarina Karničar[3,4], Aleksandra Usenik[3,4], Nataša Lindič[3,4], Andreja Sekirnik[3,4], Hideaki Tsuge[5], Vito Turk[3], Henry N Chapman[1,6,7], Winfried Hinrichs[8], Gregor Ebert[2], Dušan Turk[3,4], Alke Meents[1]

[1] Center for Free-Electron Laser Science CFEL, Deutsches Elektronen-Synchrotron DESY, Notkestraße 85, 22607 Hamburg, Germany

[2] Institute of Virology, Helmholtz Munich, Trogerstrasse 30, 81675 Munich, Germany

[3] Department of Biochemistry and Molecular and Structural Biology, Jozef Stefan Institute, Jamova 39, 1000 Ljubljana, Slovenia

[4] Centre of Excellence for Integrated Approaches in Chemistry and Biology of Proteins, Jamova 39, 1000 Ljubljana, Slovenia

[5] Faculty of Life Sciences, Kyoto Sangyo University, Kyoto 603-8555, Japan

[6] Hamburg Centre for Ultrafast Imaging, Universität Hamburg, Luruper Chaussee 149, 22761 Hamburg, Germany

[7] Department of Physics, Universität Hamburg, Luruper Chaussee 149, 22761 Hamburg, Germany

[8] Institute of Biochemistry, Universität Greifswald, Felix-Hausdorff-Str. 4, 17489 Greifswald, Germany

*This work is dedicated to the memory of Prof. Nobuhiko Katunuma, a major figure in development of epoxysuccinyl inhibitors of cysteine cathepsins.*

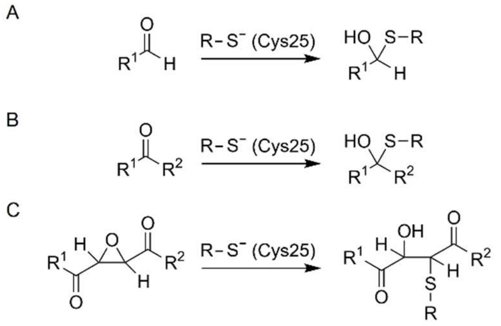


**Figure S1.** General schematic reaction of an aldehyde (**A**), a ketone (**B**) or succinyl epoxide (**C**) with the catalytic thiol of CatL. The reactions in (**A**) and (**B**) are reversible.

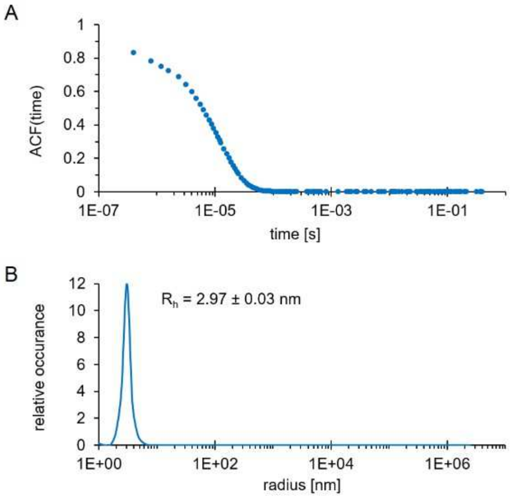


**Figure S2.** Dynamic light scattering indicating monomeric CatL in solution used for *in vitro* experiments with a hydrodynamic radius $R_h$ of 2.97 nm. The scattering data was accumulated over 100 s to obtain the auto-correlation function (ACF) shown in (**A**). The particle radius distribution shown in (**B**) was obtained using the Stokes-Einstein equation, for which the diffusion constant was determined via the CONTIN algorithm.

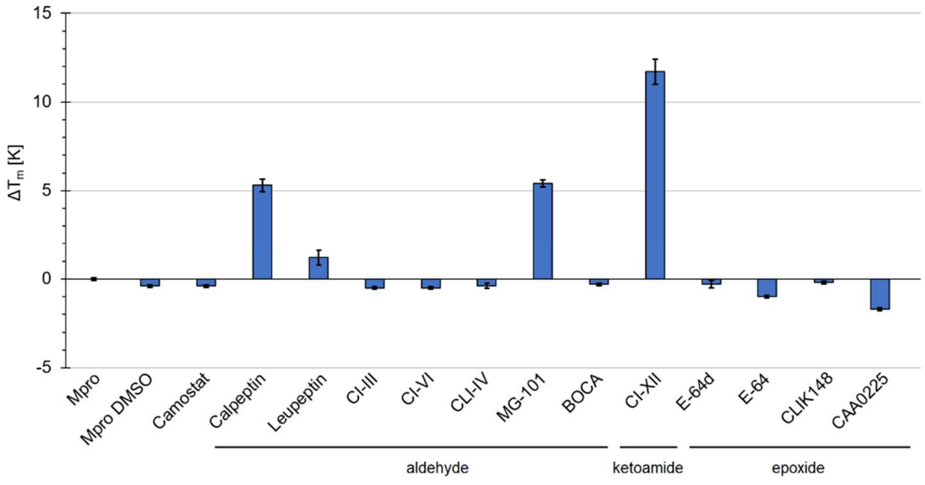


**Figure S3.** NanoDSF assay. Comparison of thermal stability of SARS-CoV-2 $M^{pro}$ as a relative measure for compound affinity. The melting temperature difference $\Delta T_m$ in the presence of a compound (relative to apo $M^{pro}$ in the absence of DMSO) is provided for a 2:1 compound to protein mixing ratio and evaluated based on a one-site binding fit using the EMBL online data-analysis platform eSPC.

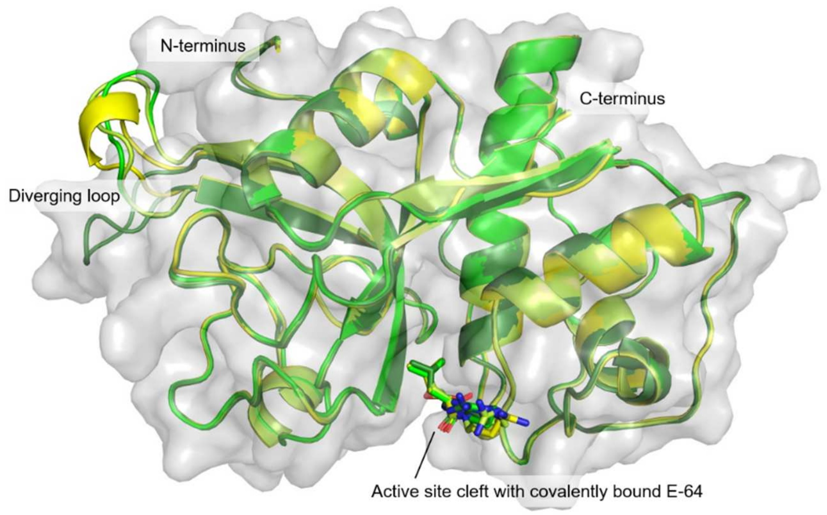


**Figure S4.** Exemplary superimposition of the four CatL chains found in the ASU (PDB 8A4V) fading from green (chain A) to yellow (chain D).

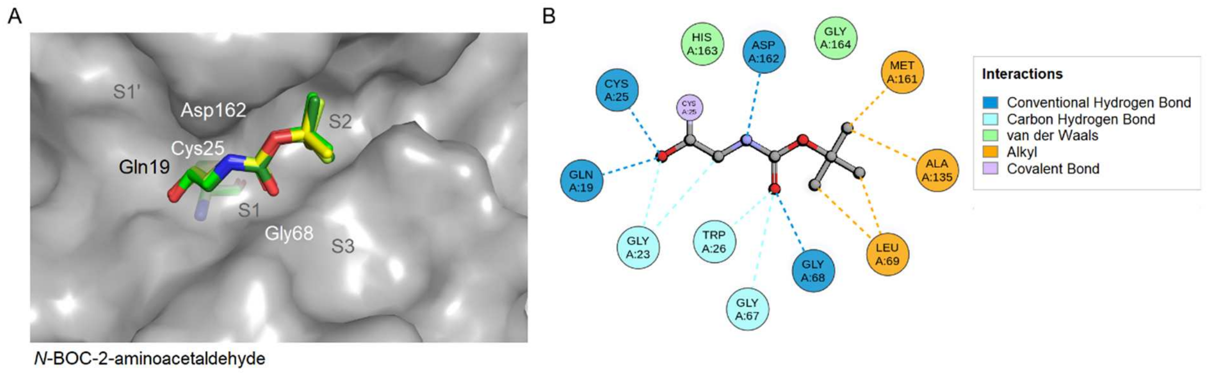


**Figure S5.** (**A**) Binding site and superposition of covalently bound *N-BOC-2-aminoacetaldehyde* from the four individual CatL molecules found in the ASU fading from green to yellow. (**B**) Two-dimensional interaction plot based on chain A according to Discovery Studio. The BOC group is located in the S2 subsite.

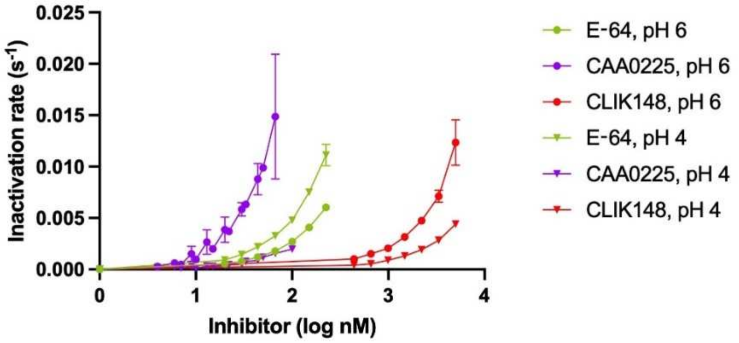


**Figure S6.** Inactivation of CatL. *In vitro* inhibition assay comparing E-64, CLIK148 and CAA0225 at pH 6 and pH 4. The inactivation rates (y-axis; $sec^{-1}$) are shown for each inhibitor concentration (x-axis; log nM). CAA0225 is shown in purple, E-64 in green and CLIK148 in red. Datapoints, obtained at pH 6 and 4 are shown as circles and triangles, respectively. The standard error boxes for each datapoint are shown.

**Table S1.** Chemical and kinetic properties of CatL inhibitors.

| Inhibitor | MW | Warhead | Binding mode | PubChem CID | $K_i$/$IC_{50}$ for CatL [nM] | $\Delta T_m$ [K] |
|---|---|---|---|---|---|---|
| **CI-III** ($C_{22}H_{26}N_2O_4$) | 382.5 | Aldehyde | Covalent | 72430 | -/- | 12.1 ± 1.1 |
| **CI-VI** ($C_{17}H_{25}FN_2O_4S$) | 372.5 | Aldehyde | Covalent | 9885817 | -/1.6 [1] | 15.2 ± 0.4 |
| **CLI-IV** ($C_{27}H_{29}N_3O_4S$) | 491.6 | Aldehyde | Covalent | 16760028 | -/1.9 [2] | 15.7 ± 0.1 |
| **MG-101** ($C_{20}H_{37}N_3O_4$) | 383.5 | Aldehyde | Covalent | 443118 | 0.5 [3]/5.8 [4] | 17.3 ± 0.6 |
| **MG-132** ($C_{26}H_{41}N_3O_5$) | 475.6 | Aldehyde | Covalent | 462382 | -/12.3 [4] | 16.6 ± 0.1 |
| **BOCA** ($C_7H_{13}NO_3$) | 159.2 | Aldehyde | Covalent | 4247255 | -/- | -0.3 ± 0.1 |
| **Calpeptin** ($C_{20}H_{30}N_2O_4$) | 362.5 | Aldehyde | Covalent | 73364 | 0.13 [5]/44.0 [4] | 18.7 ± 0.1 |
| **Leupeptin** ($C_{20}H_{38}N_6O_4$) | 426.6 | Aldehyde | Covalent | 72429 | -/0.3 [6] | - |
| **CI-XII** ($C_{26}H_{34}N_4O_5$) | 482.6 | α-Ketoamide | Covalent | 16760340 | -/1.6 [7] | 18.5 ± 0.4 |
| **13b** ($C_{31}H_{39}N_5O_7$) | 593.7 | α-Ketoamide | Covalent | 146026181 | -/- | - |
| **E-64d** ($C_{17}H_{30}N_2O_5$) | 342.4 | Epoxide | Covalent, irreversible | 65663 | 3.5 [8]*/- | 4.5 ± 0.1 |
| **E-64** ($C_{15}H_{27}N_5O_5$) | 357.1 | Epoxide | Covalent, irreversible | 123985 | -/5.5 [7] | 4.1 ± 0.4 |
| **CLIK148** ($C_{22}H_{26}N_4O_4$) | 410.5 | Epoxide | Covalent, irreversible | - (PubChem SID: 152186753) | -/<100 [9] | 13.8 ± 0.1 |
| **CAA0225** ($C_{28}H_{29}N_3O_5$) | 487.5 | Epoxide | Covalent, irreversible | 50909779 | -/1.9 [10] | 6.9 ± 2.6 |
| **TC-I** ($C_{27}H_{33}N_5O_5S$) | 539.6 | Thiocarbazate | Covalent | 16725315 | -/6.9 [11] | 18.4 ± 0.1 |
| **TPCK** ($C_{17}H_{18}ClNO_3S$) | 351.8 | Chloromethyl ketone | Covalent, irreversible | 439647 | -/<1000 [12] | 4.7 ± 0.1 |

* for the hydrolysis product E-64c

**Table S2.** Crystallographic table (1/3).

| PDB ID | 8A4X | 7ZS7 | 8A4W | 8A5B | 7QKD |
|---|---|---|---|---|---|
| **Compound** | **CI-III** | **CI-VI** | **CLI-IV** | **MG-101** | **MG-132** |
| **Data collection and unit cell*** | | | | | |
| Wavelength (Å) | 1.033 | 1.033 | 1.033 | 1.033 | 1.033 |
| Space group | P1 | P1 | P1 | P1 | P1 |
| a, b, c (Å) | 56.95, 62.25, 67.11 | 57.37, 62.56, 68.15 | 57.31, 62.58, 68.16 | 57.24, 62.2, 67.24 | 57.28, 62.71, 68.07 |
| α, β, γ (°) | 105.47, 93.52, 116.07 | 105.48, 93.43, 115.52 | 105.67, 93.33, 115.3 | 105.35, 93.25, 115.81 | 105.54 93.39 115.36 |
| Resolution (Å) | 43.92-1.8 (1.864-1.8) | 44.4-1.59 (1.6-1.63) | 44.39-1.4 (1.45-1.4) | 49.25-1.8 (1.87-1.8) | 44.35-1.5 (1.554-1.5) |
| Total reflections | 265691 (25444) | 1866980 (118533) | 1917814 (84497) | 1275006 (130857) | 1286966 (110671) |
| Unique reflections | 68306 (6823) | 101066 (7218) | 146390 (9238) | 73714 (7954) | 118682 (10402) |
| $R_{meas}$ | 0.1631 (1.319) | 0.323 (1.677) | 0.2027 (2.202) | 0.321(3.572) | 0.2597 (1.275) |
| Mean I / sigma (I) | 5.90 (1.56) | 12.40 (3.21) | 11.18 (1.70) | 7.59 (3.14) | 8.94 (1.93) |
| $CC_{1/2}$ | 0.987 (0.442) | 0.997 (0.840) | 0.998 (0.454) | 0.970 (0.68) | 0.982 (0.762) |
| Completeness (%) | 94.29 (93.28) | 93.0 (89.6) | 91.91 (57.71) | 99.3 (99.1) | 91.57 (79.93) |
| Redundancy | 3.9 (3.7) | 18.5(16.4) | 13.1 (9.1) | 17.3 (16.5) | 10.8 (10.6) |
| Wilson B-factor (Å$^2$) | 26.7 | 17.3 | 14.1 | 21.3 | 16 |
| **Refinement statistics*** | | | | | |
| Resolution (Å) | 43.92-1.8 (1.864-1.8) | 44.4-1.6 (1.6-1.63) | 44.39-1.4 (1.45-1.4) | 49.25-1.8 (1.87-1.8) | 44.35-1.5 (1.554-1.5) |
| Reflections in refinement | 68268 (6815) | 99448 (9988) | 146280 (9184) | 72723 (7259) | 118542 (10397) |
| $R_{work}$ / $R_{free}$ | 0.1921 (0.2873)/ 0.2263 (0.36) | 0.1561 (0.1739)/ 0.1877 (0.2075) | 0.1452 (0.2968)/ 0.1667 (0.3172) | 0.1784 (0.223)/ 0.2245 (0.2655) | 0.1799 (0.2833)/ 0.1976 (0.3012) |
| *No. non-hydrogen atoms* | | | | | |
| Overall | 7332 | 7840 | 8372 | 7216 | 8096 |
| Protein | 6768 | 6905 | 7101 | 6825 | 6975 |
| Ligands | 222 | 161 | 215 | 143 | 189 |
| Water | 342 | 774 | 1056 | 248 | 932 |
| *B-factors* | | | | | |
| Average | 35.9 | 21.8 | 20.4 | 28.9 | 23.8 |
| Protein | 35.6 | 20.8 | 18.6 | 28.7 | 22.1 |
| Ligands | 43.5 | 32.0 | 35 | 40.0 | 47.9 |
| Water | 36.4 | 29.0 | 29.7 | 27.6 | 31.7 |
| *RMS* | | | | | |
| Bond lengths (Å) | 0.003 | 0.0127 | 0.007 | 0.008 | 0.007 |
| Bond angles (°) | 0.59 | 1.16 | 0.83 | 0.93 | 1.17 |
| *Ramachandran* | | | | | |
| Favored (%) | 97.19 | 97.59 | 97.36 | 97.11 | 97.34 |
| Allowed (%) | 2.81 | 2.41 | 2.64 | 2.89 | 2.66 |
| Outliers (%) | 0 | 0 | 0 | 0 | 0 |

*Values in parentheses refer to the outer resolution shell.

**Table S3**. Crystallographic table (2/3).

| **PDB ID** | **8B4F** | **8AHV** | **8PRX** | **7ZXA** | **8A4V** |
|---|---|---|---|---|---|
| **Compound** | **BOCA** | **CI-XII** | **13b** | **E-64d** | **E-64** |
| **Data collection and unit cell*** | | | | | |
| Wavelength (Å) | 1.033 | 1.033 | 1.033 | 1.033 | 1.033 |
| Space group | P1 | P1 | P1 | P1 | P1 |
| a, b, c (Å) | 57.21, 62.26, 67.63 | 57.06, 62.64, 67.34 | 57.09, 62.67, 67.51 | 57.07, 62.26, 67.42 | 57.35, 62.75, 68.35 |
| α, β, γ (°) | 105.44, 93.41, 115.90 | 105.42, 93.71, 115.36 | 105.45, 93.67, 115.53 | 105.27, 93.74, 115.78 | 105.63, 93.32, 115.45 |
| Resolution (Å) | 49.42-1.9 (1.95-1.9) | 44.17-1.7 (1.75-1.7) | 49.7-1.8 (1.86-1.8) | 49.38-1.6 (1.657-1.6) | 41.9-1.65 (1.71-1.65) |
| Total reflections | 654421 (50207) | 915013 (75945) | 765727 (59054) | 1089302 (105884) | 665626 (67853) |
| Unique reflections | 117311 (8921) | 80694 (6674) | 67694 (5919) | 97574 (9570) | 95327 (9473) |
| $R_{meas}$ | 0.201 (1.394) | 0.118 (1.067) | 0.157 (0.526) | 0.1433 (2.34) | 0.3356 (2.774) |
| Mean I / sigma (I) | 7.14 (1.55) | 15.46 (2.83) | 22.66 (7.3) | 11.28 (1.03) | 5.97 (1.15) |
| $CC_{1/2}$ | 0.995 (0.68) | 0.999 (0.87) | 0.999 (0.96) | 0.999 (0.52) | 0.989 (0.368) |
| Completeness (%) | 93.8 (94.9) | 92.1 (91.6) | 91.5 (85.3) | 93.63 (91.89) | 97.57 (96.99) |
| Redundancy | 5.6 (5.6) | 11.3 (11.3) | 11.3 (10) | 11.2 (11.1) | 7.0 (7.2) |
| Wilson B-factor (Å$^2$) | 19 | 19.7 | 13.5 | 21.3 | 19 |
| **Refinement statistics*** | | | | | |
| Resolution (Å) | 49.42-1.9 (1.95-1.9) | 44.17-1.7 (1.75-1.7) | 49.7-1.8 (1.86-1.8) | 49.38-1.6 (1.657-1.6) | 41.9-1.65 (1.71-1.65) |
| Reflections in refinement | 59046 (5988) | 80674 (6542) | 67693 (6278) | 97456 (9496) | 95310 (9470) |
| $R_{work}$ / $R_{free}$ | 0.1775 (0.2708)/ 0.2099 (0.3186) | 0.1684 (0.2432)/ 0.2069 (0.3327) | 0.157(0.1897)/ 0.2030(0.2434) | 0.1636 (0.304)/ 0.1937 (0.3455) | 0.1612 (0.2713)/ 0.2029 (0.2909) |
| *No. non-hydrogen atoms* | | | | | |
| Overall | 7273 | 7455 | 7714 | 7935 | 7897 |
| Protein | 6807 | 6897 | 6793 | 6995 | 6938 |
| Ligands | 164 | 223 | 355 | 293 | 157 |
| Water | 302 | 335 | 566 | 647 | 802 |
| *B-factors* | | | | | |
| Average | 28.1 | 25.7 | 17.7 | 31.3 | 25.8 |
| Protein | 27.9 | 25.3 | 16.7 | 30.6 | 24.6 |
| Ligands | 38.1 | 36.8 | 31.6 | 41.7 | 45.5 |
| Water | 28 | 25.5 | 21.2 | 35.1 | 32.2 |
| *RMS* | | | | | |
| Bond lengths (Å) | 0.008 | 0.013 | 0.012 | 0.005 | 0.011 |
| Bond angles (°) | 0.93 | 1.24 | 1.10 | 0.77 | 0.98 |
| *Ramachandran* | | | | | |
| Favored (%) | 96.88 | 96.84 | 97.11 | 97.69 | 97.71 |
| Allowed (%) | 3.12 | 3.16 | 2.89 | 2.31 | 2.29 |
| Outliers (%) | 0 | 0 | 0 | 0 | 0 |

*Values in parentheses refer to the outer resolution shell.

**Table S4**. Crystallographic table (3/3).

| **PDB ID** | **7ZVF** | **8A4U** | **8C77** | **8OFA** |
|---|---|---|---|---|
| **Compound** | **CLIK148** | **CAA0225** | **TC-I** | **TPCK** |
| **Data collection and unit cell*** | | | | |
| Wavelength (Å) | 1.033 | 1.033 | 1.033 | 1.033 |
| Space group | P1 | P1 | P1 | P1 |
| a, b, c (Å) | 57.22, 62.29, 68.03 | 57, 62.87, 65.18 | 57.08, 62.24, 67.87 | 56.96, 62.73, 67.37 |
| α, β, γ (°) | 105.528, 93.561 115.725 | 104.04, 95.83, 116.05 | 105.25, 93.71, 115.32 | 105.07, 94.09, 115.54 |
| Resolution (Å) | 44.33-1.6 (1.657-1.6) | 44.14-1.9 (1.968-1.9) | 49.57-1.7 (1.76-1.7) | 49.62-1.9 (1.95-1.9) |
| Total reflections | 1101789 (110529) | 419485 (41509) | 836978 (60014) | 434965 (32925) |
| Unique reflections | 99390 (9947) | 56804 (5668) | 81304 (8012) | 114947 (8667) |
| $R_{meas}$ | 0.17 (0.831) | 0.3567 (2.069) | 0.400 (1.682) | 0.123 (0.479) |
| Mean I / sigma (I) | 12.78 (3.19) | 5.30 (1.18) | 10.32 (1.89) | 11.27 (3.57) |
| $CC_{1/2}$ | 0.998 (0.915) | 0.983 (0.503) | 0.995 (0.759) | 0.996 (0.911) |
| Completeness (%) | 94.3 (94.8) | 94.04 (93.3) | 92.5 (91.9) | 91.7 (92.4) |
| Redundancy | 11.1 (11.1) | 7.4 (7.3) | 10.3 (7.5) | 3.8 (3.8) |
| Wilson B-factor (Å$^2$) | 23.1 | 26.1 | 19.14 | 18.04 |
| **Refinement statistics*** | | | | |
| Resolution (Å) | 44.33-1.6 (1.657-1.6) | 44.14-1.9 (1.968-1.9) | 49.57-1.7 (1.76-1.7) | 49.62-1.9 (1.95-1.9) |
| Reflections in refinement | 99346 (9964) | 56766 (5654) | 81250 (8086) | 114894 (8123) |
| $R_{work}$ / $R_{free}$ | 0.1648 (0.2063)/ 0.1892 (0.2539) | 0.1784 (0.2842)/ 0.2088 (0.2982) | 0.1586 (0.2507)/ 0.1968 (0.2868) | 0.2171 (0.2246)/ 0.1758 (0.2580) |
| *No. non-hydrogen atoms* | | | | |
| Overall | 7667 | 7337 | 7691 | 7609 |
| Protein | 6883 | 6709 | 6852 | 6760 |
| Ligands | 287 | 241 | 248 | 293 |
| Water | 497 | 387 | 591 | 556 |
| *B-factors* | | | | |
| Average | 18.5 | 34 | 25.4 | 23.1 |
| Protein | 17.7 | 33.4 | 24.5 | 22.4 |
| Ligands | 28.6 | 48 | 38.6 | 36.6 |
| Water | 22.7 | 36.6 | 30.6 | 25.2 |
| *RMS* | | | | |
| Bond lengths (Å) | 0.007 | 0.011 | 0.007 | 0.01 |
| Bond angles (°) | 1.04 | 1.15 | 0.85 | 1.008 |
| *Ramachandran* | | | | |
| Favored (%) | 97.71 | 96.88 | 98.04 | 97.42 |
| Allowed (%) | 2.29 | 3.12 | 1.96 | 2.46 |
| Outliers (%) | 0 | 0 | 0 | 0.12 |

*Values in parentheses refer to the outer resolution shell.

#### References


1. Inoue, J. *et al.* Structure−Activity Relationship Study and Drug Profile of *N* -(4-Fluorophenylsulfonyl)- L -valyl- L -leucinal (SJA6017) as a Potent Calpain Inhibitor. *J. Med. Chem.* **46**, 868–871 (2003).

2. Yasuma, T. *et al.* Synthesis of Peptide Aldehyde Derivatives as Selective Inhibitors of Human Cathepsin L and Their Inhibitory Effect on Bone Resorption. *J. Med. Chem.* **41**, 4301–4308 (1998).

3. Sasaki, T. *et al.* Inhibitory Effect of di- and Tripeptidyl Aldehydes on Calpains and Cathepsins. *J. Enzym. Inhib.* **3**, 195–201 (1990).

4. Yang, W.-L. *et al.* Potential drug discovery for COVID-19 treatment targeting Cathepsin L using a deep learning-based strategy. *Comput. Struct. Biotechnol. J.* **20**, 2442–2454 (2022).

5. Reinke, P. *et al.* Calpeptin is a potent cathepsin inhibitor and drug candidate for SARS-CoV-2 infections. (2023), doi:10.21203/rs.3.rs-2450926/v1.

6. Mason, R. W., Green, G. D. J. & Barrett, A. J. Human liver cathepsin L. *Biochem. J.* **226**, 233–241 (1985).

7. Hu, Y. *et al.* Boceprevir, Calpain Inhibitors II and XII, and GC-376 Have Broad-Spectrum Antiviral Activity against Coronaviruses. *ACS Infect. Dis.* **7**, 586–597 (2021).

8. Towatari, T. *et al.* Novel epoxysuccinyl peptides A selective inhibitor of cathepsin B, in vivo. *FEBS Lett.* **280**, 311–315 (1991).

9. Katunuma, N. *et al.* Structure based development of novel specific inhibitors for cathepsin L and cathepsin S in vitro and in vivo. *FEBS Lett.* **458**, 6–10 (1999).

10. Takahashi, K. *et al.* Characterization of CAA0225, a Novel Inhibitor Specific for Cathepsin L, as a Probe for Autophagic Proteolysis. *Biol. Pharm. Bull.* **32**, 475–479 (2009).

11. Shah, P. P. *et al.* Kinetic Characterization and Molecular Docking of a Novel, Potent, and Selective Slow-Binding Inhibitor of Human Cathepsin L. *Mol. Pharmacol.* **74**, 34–41 (2008).

12. Lee, J.-J., Chen, H.-C. & Jiang, S.-T. Purification and Characterization of Proteinases Identified as Cathepsins L and L-like (58 kDa) Proteinase from Mackerel ( *Scomber australasicus* ). *Biosci. Biotechnol. Biochem.* **57**, 1470–1476 (1993).
